# Long-Term *In Vivo* Molecular Monitoring Using Aptamer-Graphene Microtransistors

**DOI:** 10.1101/2023.10.18.562080

**Authors:** Guangfu Wu, Eric T. Zhang, Yingqi Qiang, Colin Esmonde, Xingchi Chen, Zichao Wei, Yang Song, Xincheng Zhang, Michael J. Schneider, Huijie Li, He Sun, Zhengyan Weng, Sabato Santaniello, Jie He, Rebecca Y. Lai, Yan Li, Michael R. Bruchas, Yi Zhang

**Affiliations:** Department of Biomedical Engineering, University of Connecticut, Storrs, CT 06269, USA; Institute of Materials Science, University of Connecticut, Storrs, CT 06269, USA; Department of Bioengineering, University of Washington, Seattle, WA 98195, USA; Department of Chemical and Biomedical Engineering, FAMU-FSU College of Engineering, Florida State University, Tallahassee, FL 32306, USA; Department of Chemistry, University of Connecticut, Storrs, CT 06269, USA; Department of Chemistry, University of Nebraska-Lincoln, Lincoln, NE 68588, USA; Department of Anesthesiology and Pain Medicine, University of Washington, Seattle, WA 98195, USA; Center for Neurobiology of Addiction, Pain, and Emotion, University of Washington, Seattle, WA 98195, USA; Department of Pharmacology, University of Washington, Seattle, WA 98195, USA

**Author notes:** G.W., E.T.Z., and Y.Q. contributed equally to this work.

## Abstract

Long-term, real-time molecular monitoring in complex biological environments is critical for our ability to understand, prevent, diagnose, and manage human diseases. Aptamer-based electrochemical biosensors possess the promise due to their generalizability and a high degree of selectivity. Nevertheless, the operation of existing aptamer-based biosensors *in vivo* is limited to a few hours. Here, we report a first-generation long-term *in vivo* molecular monitoring platform, named aptamer-graphene microtransistors (AGMs). The AGM incorporates a layer of pyrene- (polyethylene glycol)5-alcohol and DNase inhibitor-doped polyacrylamide hydrogel coating to reduce biofouling and aptamer degradation. As a demonstration of function and generalizability, the AGM achieves the detection of biomolecules such as dopamine and serotonin in undiluted whole blood at 37 °C for 11 days. Furthermore, the AGM successfully captures optically evoked dopamine release *in vivo* in mice for over one week and demonstrates the capability to monitor behaviorally-induced endogenous dopamine release even after eight days of implantation in freely moving mice. The results reported in this work establish the potential for chronic aptamer-based molecular monitoring platforms, and thus serve as a new benchmark for molecular monitoring using aptamer-based technology.

## Introduction

The ability for real-time monitoring of specific molecules in complex biological environments found *in vivo* has been reported to play an important role in the diagnosis and management of human diseases such as mental health, diabetes, cardiac diseases, acute kidney disease, and cancer^1–5^. For example, the continuous glucose meter (CGM), with the capability of real-time glucose monitoring for one to two weeks in the skin interstitial fluid, has revolutionized the effective management of diabetes^6^. However, the CGM represents the only commercially available biosensor for real-time molecular monitoring *in vivo.* The enzymatic sensing mechanism of CGM is not generalizable to other important biomarkers, like drugs^7–10^ and hormones^11–13^, due to the limited number of available enzymes for targeted analytes. To date, the design of biosensors for real-time molecular monitoring beyond the CGM has faced many of the following technical challenges: (1) the biosensing platform must be generalizable, thereby making it easy to adapt the platform to new targets, (2) it must have sufficient sensitivity and selectivity both for the low (nM) concentrations of biomarkers and be robust to interference from other molecules with similar structures, and (3) it must be resistant to biofoulings and performance degradation for long-term and real-time monitoring.

Electrochemical aptamer-based (EAB) biosensors have demonstrated the potential to address the abovementioned challenges for real-time molecular monitoring in clinically relevant *in vivo* environments^14–18^. Generally, EAB sensors consist of single-stranded DNA or RNA aptamers functionalized on the surface of a gold working electrode^17^. Redox reporters, like methylene blue or ferrocene, are typically functionalized to one end of the aptamers, allowing for charge transfer upon the conformational change of aptamers caused by the binding of targets^19^. More specifically, the binding and dissociation of target analytes to aptamers cause the conformational rearrangement of aptamers, which changes the electron transfer rate between redox reporters and the working electrode and results in a measurable current change. This current change can be monitored by several electrochemical methods, like square-wave voltammetry and cyclic voltammetry^20^. EAB sensors have been widely used for real-time monitoring of multiple therapeutic drugs (kanamycin^21, 22^, vancomycin^3^, ampicillin^23^, and doxorubicin^24^) and metabolites (ATP and phenylalinine^25, 26^) *in vivo* in small animal models. Nevertheless, the EAB sensors suffer from (1) limited *in vivo* measurement time (a few hours, **Supplementary Table 1**) and (2) difficulty in miniaturizing the EAB sensors down to micrometer scale (to be comparable with the CGM) considering the tradeoff between the magnitude of measurable current and the microscale sensor dimensions^14, 27, 28^.

Unlike simple electrode-based electrochemical transducers, transistors can simultaneously transduce and amplify the sensor signal, thereby enabling the miniaturization of aptamer-based electrochemical biosensors without sacrificing the signal-to-noise ratio^29^. Among these, graphene field-effect transistor (gFET) has emerged as a high-performance platform for electrochemical biosensing due to the high carrier mobility of graphene and its atomic layer thickness^30^, which enables the detection with high sensitivity. A graphene field-effect transistor biosensor is typically a three-terminal electronic device, in which source and drain terminals are bridged by sensing material, and current flow between source and drain electrodes is modulated by a gate terminal^31^. When the gate voltage is applied onto the surface of an gFET device through an electrolyte, positive and negative charges are separated and form an electrostatic double layer, serving as an insulator to tune the conductivity of the device^32–34^. Transistor-based aptamer sensors have been used as the new toolkits for real-time molecular monitoring *in vivo*^35, 36^. Nevertheless, like EAB sensors, these sensors can survive for only several hours in complex *in vivo* biological environments because of sensor degradation caused by biofouling, which impedes chronic performance^35, 37^.

Here, we report aptamer-graphene microtransistors (AGMs) that incorporate a layer of pyrene- (polyethylene glycol)5-alcohol (pyrene-PEG5-alcohol) and DNase inhibitors doped polyacrylamide hydrogel for long-term molecular monitoring in undiluted whole blood samples and *in vivo* environments. *In vitro* demonstrations in undiluted whole blood at 37 °C showed the detection of a physiologically relevant range of dopamine and serotonin over 11 days (retaining 50% of the original response). These surface-coated AGMs exhibited the capability to monitor optogenetically evoked dopamine release *in vivo* in mice over one week. In freely moving mice, they successfully monitored endogenous dopamine release induced by animal behaviors such as the return of righting reflex (RORR) and sucrose consumption even after eight days of implantation. The development of this type of long-term aptamer biosensor for *in vivo* monoamine monitoring will serve as a benchmark and template for developing aptamer-based biosensors for long-term monitoring of a broad range of molecules in a host of biological environments.

## Results

### Design and fabrication of AGMs

As shown in **Figs. 1a and b**, we designed an implantable filament probe (450 μm wide, 81.8 μm thick, and 6.2 mm long) that consists of four AGMs with a size of 50 µm × 50 µm. The AGM fabrication is based on photolithography and sputtering deposition protocols to define the source, drain, and gate contact pads on the polyimide substrates (thickness: 81 µm), followed by transferring patterned graphene onto the source-drain contact pads, as described in our previous studies^35, 36, 38^. Here, an Ag/AgCl electrode, widely used for acute and chronic *in vivo* studies^16, 39–42^, was fabricated on the polyimide substrate as the gate electrode. The sensor surface was then encapsulated with SU-8 layer (thickness: 800 nm) to define the active graphene sensing area of 50 μm × 50 μm, and to prevent the penetration of biofluids. The aptamers were functionalized on the surface of the graphene microtransistors. Real-time dopamine monitoring is achieved through the reversible conformational rearrangement of negatively charged phosphate backbones of aptamers functionalized on the graphene microtransistors in the presence of the target dopamine (**Fig. 1c**), thereby causing an increase in the number of hole carriers and generating a measurable shift of the transfer curve of the AGM (**Fig. 1d**). This shift is proportional to the target concentration^36^. In this study, the sensor response is defined as the percentage change of the Dirac point voltage, V_DP_ (the voltage at the lowest source-drain current in the transfer curve), caused by the presence of dopamine.

Sensor response (%) = (ΔV_DP_/V_0_) × 100

where ΔV_DP_ and V_0_ are the change of the Dirac point voltage in the presence of dopamine and the initial voltage without dopamine, respectively.

**Figure 1.**
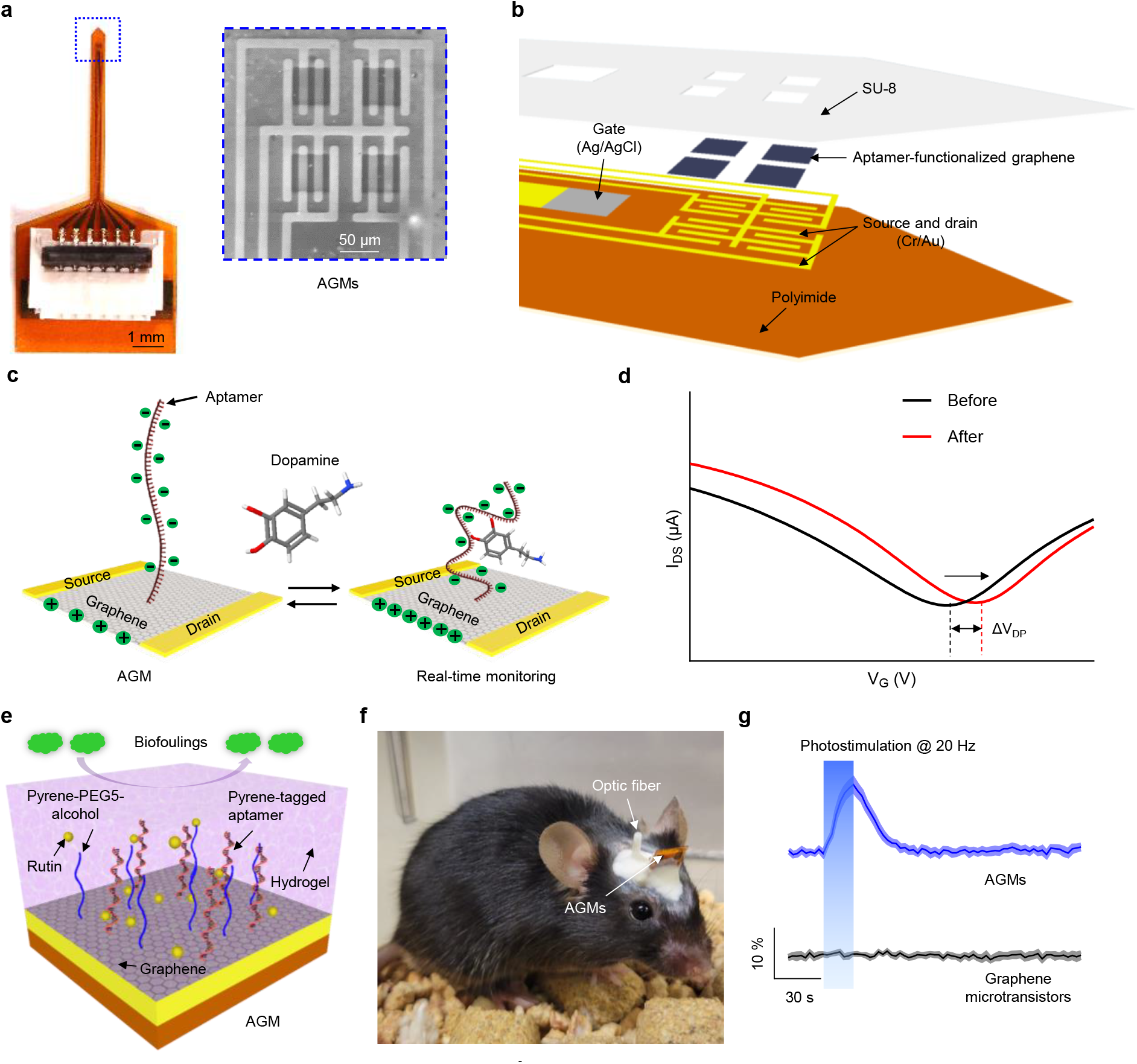
Design and features of implantable AGMs with surface coatings for long-term *in vivo* molecular monitoring. (**a**) Optical image of the overall configuration of the probe. Inset: Scanning electron microscope (SEM) image of the tip end of device that contains four AGMs. (**b**) Layered schematic illustration of the tip end of the sensor, consisting of four AGMs and an onboard Ag/AgCl gate electrode. (**c**) Working principle of the AGMs for real-time dopamine monitoring. The reversible conformational switch of aptamers in the presence of targets increases the number of hole carriers in the graphene channel surface, leading to a measurable shift of transfer curves. (**d**) Representative transfer curve shift of the AGM before and after exposure to the target. (**e**) Schematic illustration of surface coatings with a monolayer of pyrene-PEG5-alcohol and DNase inhibitors-doped polyacrylamide hydrogel on the AGMs. (**f**) Representative image of a mouse following the implantation of the AGMs. (**g**) Representative real-time monitoring of *in vivo* dopamine release in nucleus accumbens shell (NAcSh) of mice induced by photostimulation (20 Hz, 5 ms pulse width). The shading area represents ± SEM from seven samples.

The sensor could lose functionality in complex biological environments due to: **1**) the loss of surface assembled monolayer (aptamers); **2**) biofouling of the sensor surfaces due to non-specific adsorptions**; 3**) the degradation of aptamer from endogenous nucleases in *in vivo* environments. To tackle these issues and enable long-term molecular monitoring, the sensor surface was first functionalized with a dopamine aptamer, followed by surface passivation with pyrene-PEG5- alcohol and DNase inhibitor-doped anti-biofouling hydrogel surface coatings (thickness: ∼10 µm) (**Fig. 1e and Supplementary Fig. 1**). **Fig. 1f** shows a mouse implanted with the surface-coated AGM in the nucleus accumbens shell (NAcSh) of a mouse brain following one week of implantation. **Fig. 1g** demonstrates the capability of real-time *in vivo* dopamine monitoring during photostimulation (20 Hz, 5 ms pulse width) for surface-coated AGMs, whereas the control graphene microtransistor channel (without aptamer functionalization) shows negligible signal response during photostimulation.

### Aptamer monolayer functionalized with non-covalent π-π stacking is stable

To examine the potential loss of functionalized aptamers on the graphene surface, we investigated the stability of aptamer monolayers based on two widely used functionalization methods, covalent electrografting and non-covalent π-π stacking (**Fig. 2a**). For the covalent method, the graphene surface was first functionalized with 4-aminophenyl (Ph-NH_2_) group through the electrochemical grating method. The electrografted NH_2_ group served as the linker to anchor dopamine aptamer through EDC/NHS reaction, as reported in our previous work^36^. For the non-covalent method, pyrene-tagged dopamine aptamer was assembled onto the graphene surface via the π-π interaction between the pyrene group and graphene. Successful surface functionalization based on these two methods was verified via Raman spectroscopy (**Figs. 2b and c**). A clear D peak was observed for the electrografted graphene sample, which is attributed to the conversion of sp^2^ hybridized carbons to sp^3^ ones. Additionally, an obvious G’ peak at ∼1623 cm^-1^ was obtained in the non-covalent case, resulting from the pyrene molecule attached to the graphene surface via π-π stacking^43^. The sensing performance of the AGMs prepared using these two different methods was determined by exposing sensors to biologically relevant dopamine concentrations (1 nM, 100 nM, and 10 μM). As shown in **Figs. 2d and e**, the transfer curves of AGMs shifted rightward when dopamine solution was added, and a higher dopamine concentration caused a larger shift of the transfer curves, suggesting that both methods are able to successfully functionalize aptamers on the graphene microtransistors.

**Figure 2.**
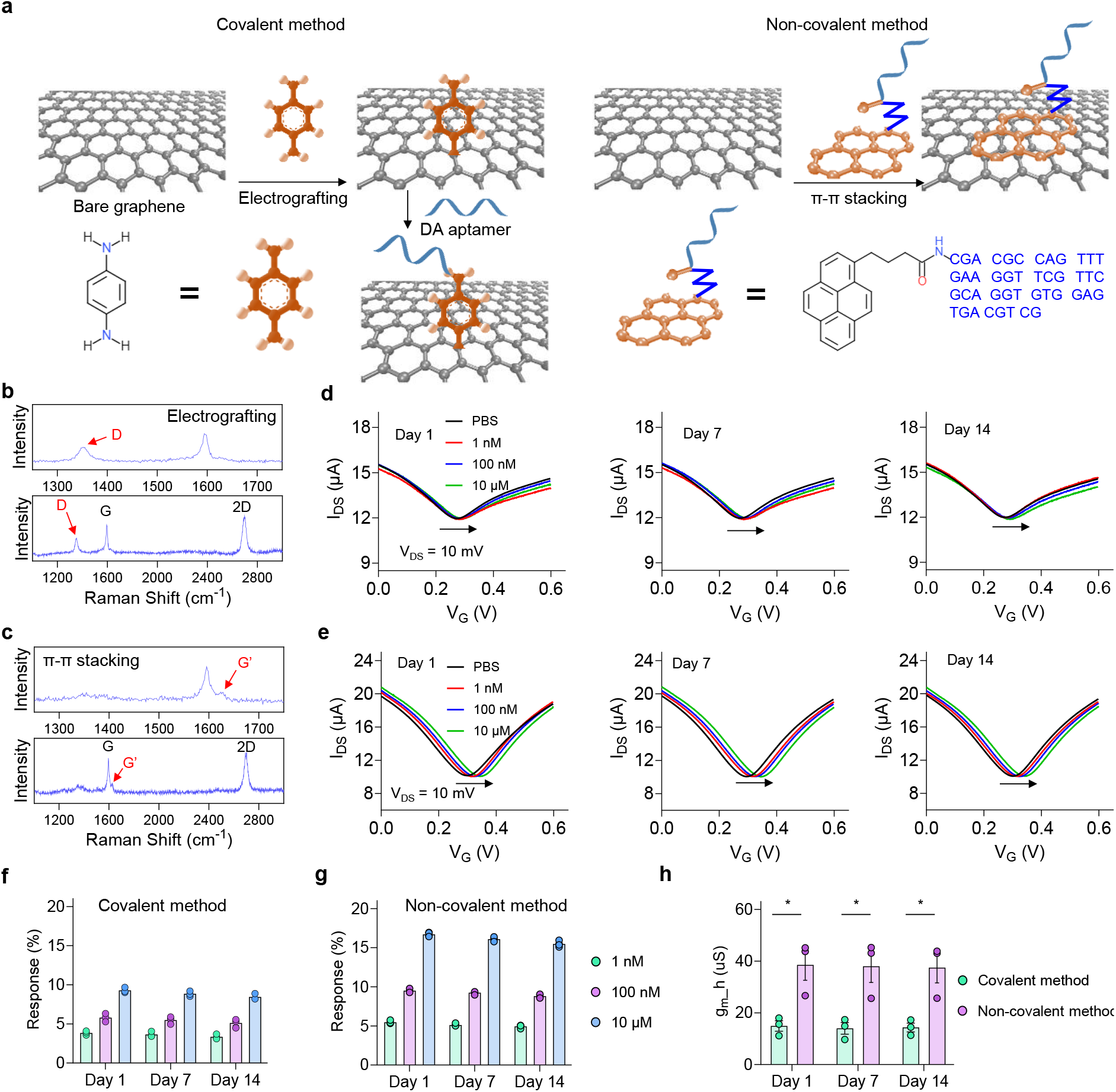
Characterization of aptamer monolayers functionalized with covalent electrografting and non-covalent π-π stacking methods. (**a**) Schematic illustration of functionalizing aptamers on the graphene surface with electrochemical grafting (left) and π-π stacking (right). (**b**) Raman spectrum of graphene after electrografting with p-phenylenediamine. (**c**) Raman spectrum of graphene after the functionalization with pyrene-tagged aptamers using π- π stacking. (**d**) The response of sensors using the covalent functionalization method after incubating in 1× PBS at 37 °C for two weeks. (**e**) The response of sensors using the non-covalent π-π stacking method after incubating in 1× PBS at 37 °C for two weeks. Dopamine concentration- dependent response of AGMs prepared based on (**f**) covalent electrografting method and (**g**) non- covalent π-π stacking method across day 1, day 7, and day 14. (**h**) The transconductance (g_m_h_) of AGMs based on different aptamer functionalization methods. n = 3; \**P* < 0.05. All data are represented as means ± SEM.

We then investigated the stability of the aptamers functionalized on the sensor surface using these two methods. We incubated sensors in 1× PBS at 37 °C for two weeks, and then measured their responses to different concentrations of dopamine (**Figs. 2d and e**). Dopamine concentration-dependent electrical responses for covalent and non-covalent functionalization are summarized in **Figs. 2f and g**. Notably, minimal change of the response was observed after incubating in 1× PBS at 37 °C for two weeks for functionalized sensors based on either covalent or non-covalent method, indicating the long-term stability of the functionalized aptamers in 1× PBS. However, the overall dopamine concentration-dependent responses for sensors prepared based on the non-covalent method are much higher than those based on the covalent method (**Figs. 2f and g**). The non-covalent method preserves the intrinsic high carrier mobility of graphene microtransistors, while the covalent method lowers carrier mobility due to the conversion of sp^2^ hybridized carbons into sp^3^ hybridized ones, locally disrupting the conjugation of graphene and lowering sensor transconductance (g_m_h_) (**Fig. 2h**). Thus, a smaller sensor response is observed for AGMs prepared by the covalent method. Considering the high sensor response, simple functionalization steps, and the stable response over two weeks in 1× PBS at 37 °C, the non-covalent method is selected to functionalize aptamers onto the graphene microtransistor surface.

### Pyrene-PEG5-alcohol passivation enables long-term operation in protein-rich aCSF and rat CSF

In addition to the stability of the assembled aptamer monolayer, biofouling of the sensor surface due to non-specific adsorption represents another potential challenge for a sensor to be used in complex biological environments. We next passivated the AGMs with pyrene-PEG5- alcohol, a gold-standard surface passivation molecule, of which the pyrene molecule is attached to the graphene surface, and the PEG5-alcohol is surrounded by water molecules to serve as the hydration layer to preclude the attachment of lipophilic molecules such as proteins, enzymes, and antibodies^44–46^. We then investigated the longevity of the sensors in protein-rich artificial cerebrospinal fluid (aCSF with 5 mg/mL bovine serum albumin, BSA, to mimic a protein-rich bodily fluid^47^), rat cerebrospinal fluid (CSF), and undiluted rat whole blood samples at 37 °C. More specifically, the sensor surface was first functionalized with the dopamine aptamers, followed by the functionalization with pyrene-PEG5-alcohol, as shown in **Fig. 3a**. To study the longevity of the sensors in protein-rich biological environments, we incubated the sensors in aCSF with 5 mg/mL BSA for one week (**Fig. 3b** and **Supplementary Figs. 2a-d**). As expected, we observed a reduction of response when the sensors without a pyrene-PEG5-alcohol passivation were exposed to a high concentration of 10 µM dopamine solution, which is attributed to the reduction of available aptamers caused by non-specific adsorption of BSA. In contrast, the sensor coated with pyrene-PEG5-alcohol maintains the original performance towards physiologically relevant dopamine solutions (1 nM, 100 nM, and 10 µM) over one week, indicating that pyrene-PEG5- alcohol can be used as an effective passivation layer to reduce protein adsorption in protein-rich aCSF for one week. The effective surface passivation using pyrene-PEG5-alcohol is further supported by incubating the sensors with and without the passivation in rat CSF at 37 °C for one week (**Fig. 3c, Supplementary Figs. 2e-h**). The pyrene-PEG5-alcohol molecules form a stable hydration layer on the graphene surface^44, 48^, resulting in the reduction of non-specific adsorption and sensor performance degradation. Encouraged by the performance of the AGMs with pyrene- PEG5-alcohol in protein-rich aCSF (with 5 mg/mL BSA) and rat CSF, we finally investigated their long-term stability in a more challenging undiluted rat whole blood sample (BioIVT). As expected, the sensors showed a significant response reduction as soon as 10 hours after incubating in undiluted rat whole blood samples, though the pyrene-PEG5-alcohol protected AGMs showed higher responses than those without the pyrene-PEG5-alcohol coating (**Fig. 3d and Supplementary Figs. 2i-l**). These results indicate that pyrene-PEG5-alcohol enables sustained operation in protein-rich aCSF and rat CSF but is not effective in minimizing performance degradation in undiluted whole blood samples.

**Figure 3.**
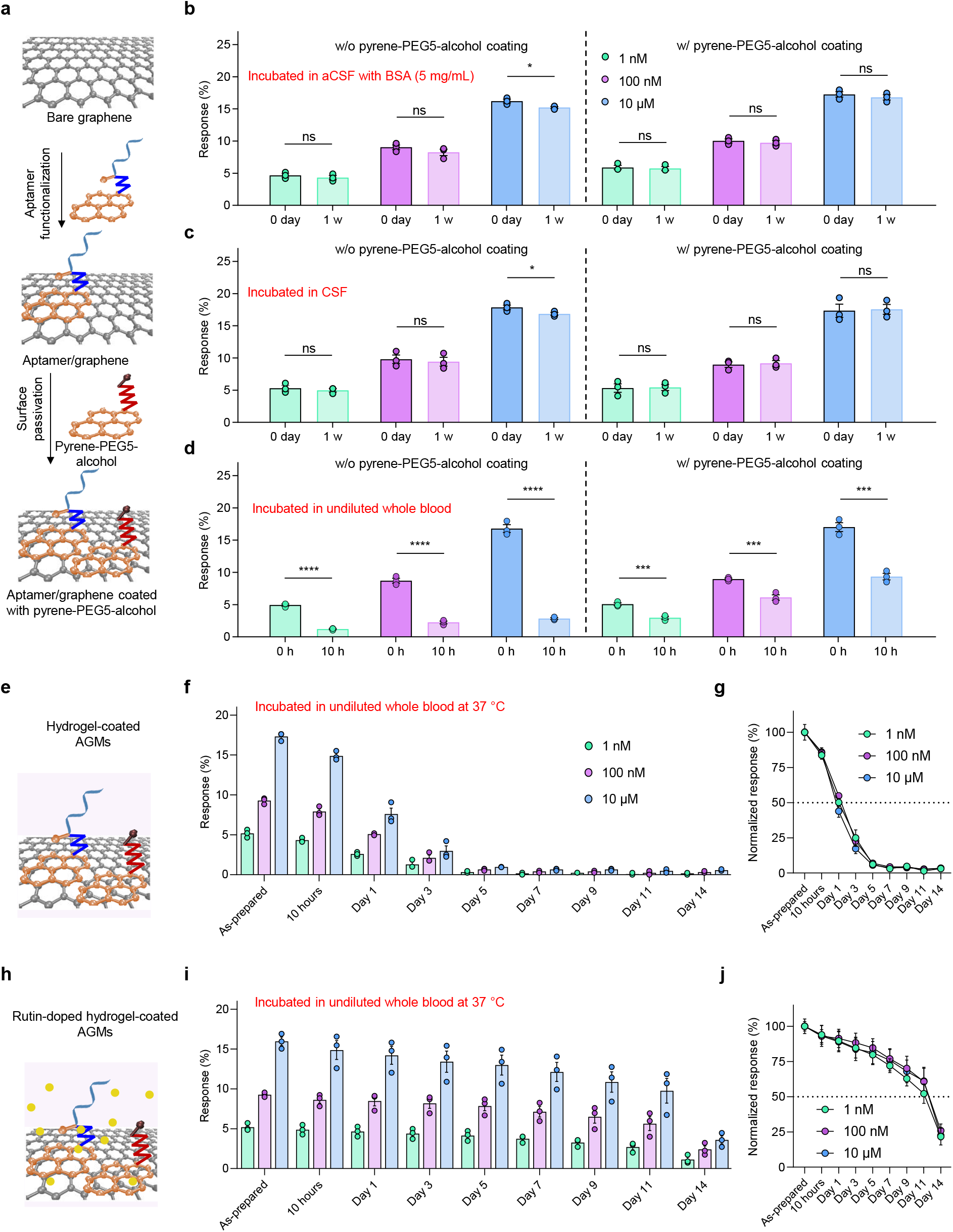
Long-term stability of AGMs with various surface coatings in aCSF, CSF, and undiluted whole blood at 37 °C. (**a**) Schematic illustration of surface functionalization with pyrene-tagged aptamer and surface passivation with a monolayer of pyrene-PEG5-alcohol. (**b**) Dopamine concentration-dependent response of AGMs with and without the pyrene-PEG5- alcohol passivation incubated in (**b**) protein-rich aCSF solution (5 mg/mL BSA protein), (**c**) rat CSF, and (**d**) undiluted whole blood samples at 37 °C. n = 3; \*\*\*\**P* < 0.0001, \*\*\**P* < 0.001, \**P* < 0.05. (**e**) Schematic illustration of pyrene-PEG5-alcohol and polyacrylamide hydrogel-coated AGMs. (**f**) The response of AGMs coated with pyrene-PEG5-alcohol and polyacrylamide hydrogel incubated in undiluted rat blood samples at 37 °C. n = 3. (**g**) Normalized sensor response after incubating in undiluted whole blood at different time points (10 hours, 1 day, 3 days, 5 days, 7 days, 9 days, 11 days, and 14 days) for pyrene-PEG5-alcohol and polyacrylamide hydrogel-coated AGMs. (**h**) Schematic illustration of pyrene-PEG5-alcohol and DNase inhibitor rutin doped polyacrylamide hydrogel-coated AGMs. (**i**) The response of AGMs coated with pyrene-PEG5- alcohol and rutin-doped polyacrylamide hydrogel incubated in undiluted rat blood samples at 37 °C. n = 3. (**j**) Normalized sensor response after incubating in undiluted whole blood at different time points (10 hours, 1 day, 3 days, 5 days, 7 days, 9 days, 11 days, and 14 days) for pyrene- PEG5-alcohol and rutin-doped polyacrylamide hydrogel-coated AGMs. All data are represented as means ± SEM.

### DNase inhibitor-doped polyacrylamide hydrogel coatings enable long-term operation in undiluted whole blood

Polyacrylamide hydrogels based on a 50:50 copolymer of hydroxyethylacrylamide and diethylacrylamide have been demonstrated to have excellent anti-biofouling property^39^. Inspired by this study, the pyrene-PEG5-alcohol passivated AGMs were further coated with a thin polyacrylamide hydrogel layer (**Fig. 3e**). The AGM with polyacrylamide hydrogel coating maintains the functionality for a few hours in undiluted rat whole blood samples. However, the overall sensor response showed ∼ 50% decrease after incubating in undiluted rat blood samples for one day (**Figs. 3f and g, Supplementary Fig. 3**). The sensors completely lost functionality and responses after three days. We hypothesized that endogenous nucleases found in undiluted whole blood samples could cause the degradation of aptamers, resulting in the loss of sensor performance.

To test our hypothesis, a DNase inhibitor, rutin^49^, was doped into the polyacrylamide hydrogel layer, resulting in a final rutin concentration of 150 μM. The pyrene-PEG5-alcohol passivated AGM was then coated with DNase inhibitors-doped polyacrylamide hydrogel (**Fig. 3h**). The transfer curves of AGMs with pyrene-PEG5-alcohol and rutin-doped polyacrylamide hydrogel were monitored when the sensors were exposed to dopamine solutions with different concentrations (**Supplementary Fig. 4**). The sensor showed dopamine concentration-dependent responses in undiluted whole blood for about two weeks (**Fig. 3i**). **Fig. 3j** shows that the sensors maintained 50% of the original signal after incubating in undiluted blood samples at 37 °C for 11 days, which is 11 times longer than that of the sensors without the addition of rutin. In addition, the incubation of AGMs with surface coating in undiluted whole blood for two weeks showed minimal influence on the transconductance for both hydrogel-coated AGMs and rutin-doped hydrogel-coated AGMs (**Supplementary** Fig. 5), in which over 95% and 90% of the original transconductance were obtained for DNase-doped hydrogel-coated AGMs and hydrogel-coated AGMs, respectively.

We then wonder whether the DNase inhibitor-doped polyacrylamide hydrogel coatings are generalizable for enabling the long-term operation of AGMs to target biomarkers beyond dopamine in undiluted whole blood. To this end, we selected AGM serotonin sensors as a model system for evaluation. Specifically, serotonin-specific aptamers were functionalized onto the graphene surface via the pyrene group, followed by surface passivation with the abovementioned pyrene-PEG5-alcohol and rutin-doped polyacrylamide hydrogel. The AGM serotonin sensors were then incubated in undiluted rat blood at 37 °C for two weeks, and the sensor response to biologically relevant concentrations of serotonin (0.1 nM, 10 nM, and 1 μM) was assessed at day 1, day 3, day 5, day 7, day 9, day 11, and day 14. Consistent with the results from AGM dopamine sensors, the serotonin sensors showed concentration-dependent responses in undiluted whole blood for about two weeks, maintaining 50% of the original signal after incubating in undiluted blood samples at 37 °C for 11 days (**Supplementary Fig. 6**). Collectively, these findings indicate the generalizability of the DNase inhibitors-doped polyacrylamide hydrogel coatings for use in aptamer-based biosensors.

### Biocompatibility of DNase inhibitor-doped polyacrylamide hydrogel coatings

Next, we tested the biocompatibility of the enhanced-longevity surface coatings, as this is crucial for their eventual *in vivo* operation. We investigated the biocompatibility of DNase inhibitor-doped polyacrylamide hydrogel coatings by coculturing them with forebrain organoids. MTT (3-(4,5-Dimethylthiazol-2-yl)-2,5-diphenyltetrazolium bromide), which indicates metabolic activity, is used to evaluate cellular activities after coculturing with the AGMs with the surface coatings. MTT absorbance shows that cell metabolic activities over 1, 4, 8, 12, and 16 days for AGMs and AGMs with surface coatings are comparable to those of the cell-only group (**Fig. 4a**). With increasing culture time, rutin-doped hydrogel-coated AGMs demonstrated no significant decrease in absorbance when compared to the cell-only group. These results establish the biocompatibility of DNase inhibitor-doped polyacrylamide hydrogel coatings.

**Figure 4.**
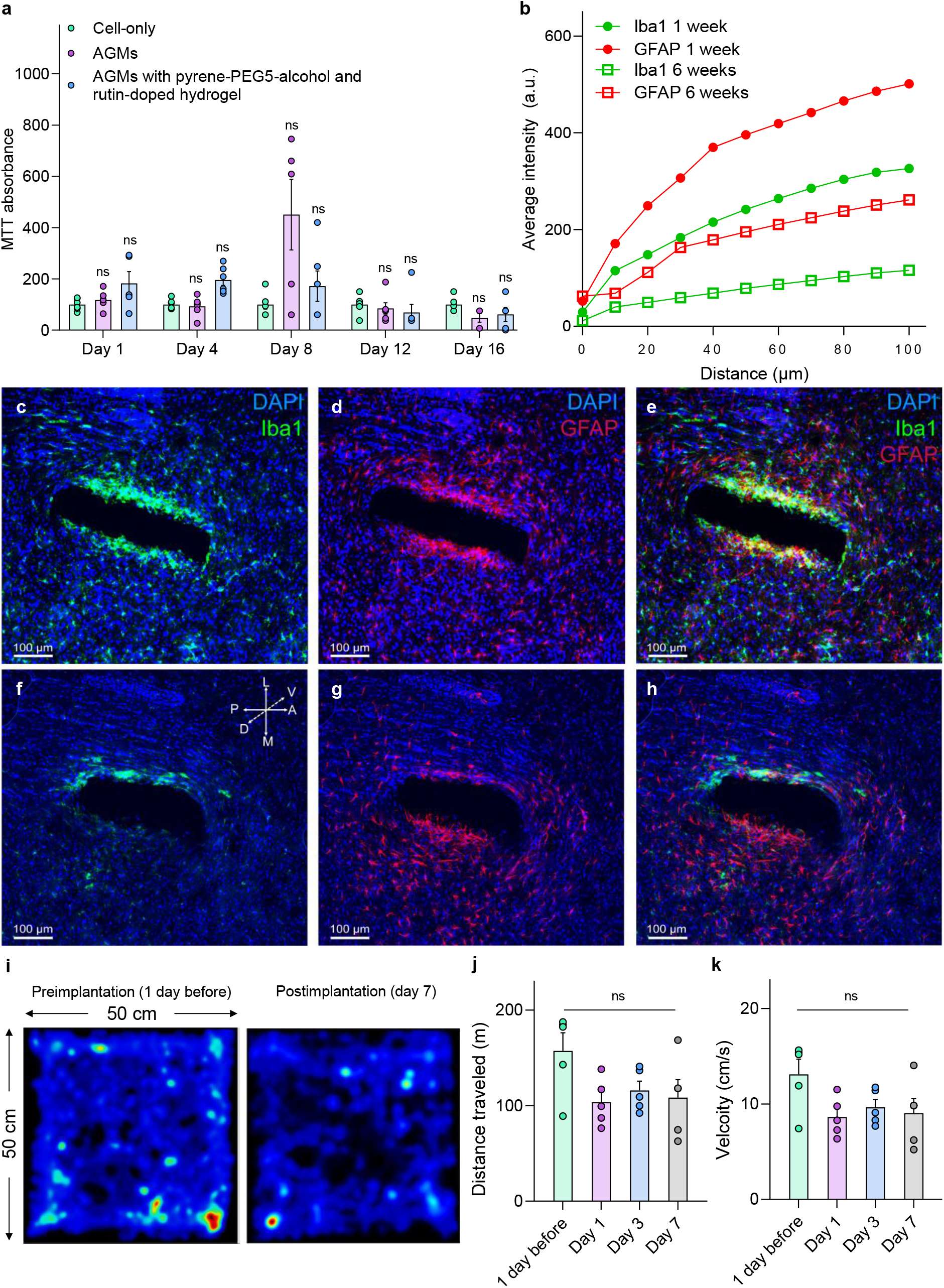
Biocompatibility of DNase inhibitor-doped polyacrylamide hydrogel coatings and the impact of sensor implantation on brain tissue and animal behaviors. (**a**) MTT absorbance of induced neural progenitor cells (iNPCs) cocultured with AGMs with different coatings. ns refers to not significant. (**b**) Representative changes in intensity within region of interest (ROI) as the distance from sensors increases for both Iba1 (green) and GFAP (red) at 1 week (closed circle) and 6 weeks (open square). (**c-h**) Confocal fluorescence images of horizontal brain slices show immunohistochemical staining for DAPI (blue), astrocytes (GFAP, red), and activated microglia (Iba1, green) and overall lesion from sensor implantation after 1 week (**c-e**) and 6 weeks (**f-h**). All histological and confocal settings were kept consistent across groups, 20× (Scale bars, 100 μm). (**i**) Animal motion heatmap for preimplantation (1 day before implantation) and postimplantation (after 7 days). (**j**) Total traveling distance and (**k**) velocity for mice in an open-field assay at four different stages: 1 day before implantation, and 1, 3, and 7 days postimplantation. n = 5; ns indicates that the difference of the means is not significant at the 0.05 level. All data are represented as means ± SEM.

### *In vivo* immunohistochemistry

To further evaluate the biocompatibility of the AGMs coated with DNase inhibitor-doped polyacrylamide hydrogel, we investigated inflammatory responses via measuring device-induced gliosis, a reactive response to brain injury, within the deep brain AGM sensor implants. The sensors were implanted into the NAcSh of mice, and the glial immune response after 1 and 6 weeks of sensor implantation surgery was evaluated by immunohistochemical staining for DAPI (blue), astrocytes (GFAP, red), and activated microglia (Iba1, green) (**Fig. 4b**). Under consistent imaging settings, the post-implantation fluorescence intensity indicates a significant decrease in the immune response over the 6-week recovery period (**Figs. 4c-h**). Moreover, the immune response and tissue lesion area induced by our sensors were equivalent to those induced by other advanced thin-film deep brainimplants^50–52^.

### Effect of sensor implantation on natural locomotor behavior

We next examined the effects of polyacrylamide hydrogel-coated AGM probe implantation on the basal anxiety state and locomotor activity of mice to determine if the implant scheme would cause any aberrant behavioral effects. We implanted the probe into the NAcSh of mice (a dopamine rich brain region)^53^ and monitored their activity in an open field test 1 day prior to implantation and 1, 3, and 7 days after implantation, by recording locomotor activity and velocity. The heatmaps for animal movement revealed comparable levels of locomotor activity between the day prior to implantation and seven days after implantation (**Fig. 4i**). Moreover, no significant differences were found over time when comparing preimplantation (1 day) and post-implantation (7 days) in both distance traveled and average velocity (**Figs. 4j and k**). In addition, mouse body weight and temperature across this time window remained stable (**Supplementary Fig. 7**). Overall, these results establish that the sensor implantation does not have significant effects on basal anxiety and locomotor activity in the mice over the prolonged duration of the implantation period.

### Long-term monitoring of optically evoked dopamine release *in vivo* over one week

Encouraged by the long-term operational characteristics from the AGMs coated with DNase inhibitor-doped polyacrylamide hydrogel in vitro, we next evaluated the capability for long-term monitoring *in vivo* using a mouse model. Cre-recombinase-dependent viral vectors containing the light-sensitive cation channel channelrhodopsin-2 (ChR2) were unilaterally injected into the ventral tegmental area (VTA) of DAT-cre mice, with a fiber optic ferrule implanted above the VTA and the AGM probe with surface coating implanted into the NAcSh (**Figs. 5a and b**). We then delivered photostimulation to the VTA (20 Hz, 5 ms pulse width), a well-established parameter for inducing phasic dopamine release^54–56^, while monitoring the resulting dopamine release in the NAcSh using the AGM by performing continuous transfer curve scanning. As shown in **Fig. 5c**, the transfer curves first shift to the rightward during photostimulation and then shift back to the left after stimulation ends. Accordingly, the sensor response shows a rapid increase during stimulation and a slow decrease following the end of stimulation (**Fig. 5d**). We then investigated the long-term monitoring of optically evoked dopamine release over 10 days (**Fig. 5e and Supplementary Fig. 8**). The AGMs with surface coatings successfully captured the optogenetically evoked dopamine release from day 1 through day 7. The peak response gradually decreased over subsequent days, and no clear response peak was obtained on day 10. In contrast, the sensors coated with pyrene-PEG5-alcohol and polyacrylamide hydrogel (no rutin) showed an *in vivo* operation lifetime of 2-3 days (**Supplementary Fig. 9**), further highlighting the importance of rutin doping to prevent *in vivo* sensor performance degradation from endogenous nucleases. **Fig. 5f** shows the sensor response before and during photostimulation, in which only the optically evoked dopamine release caused a robust and significant response.

**Figure 5.**
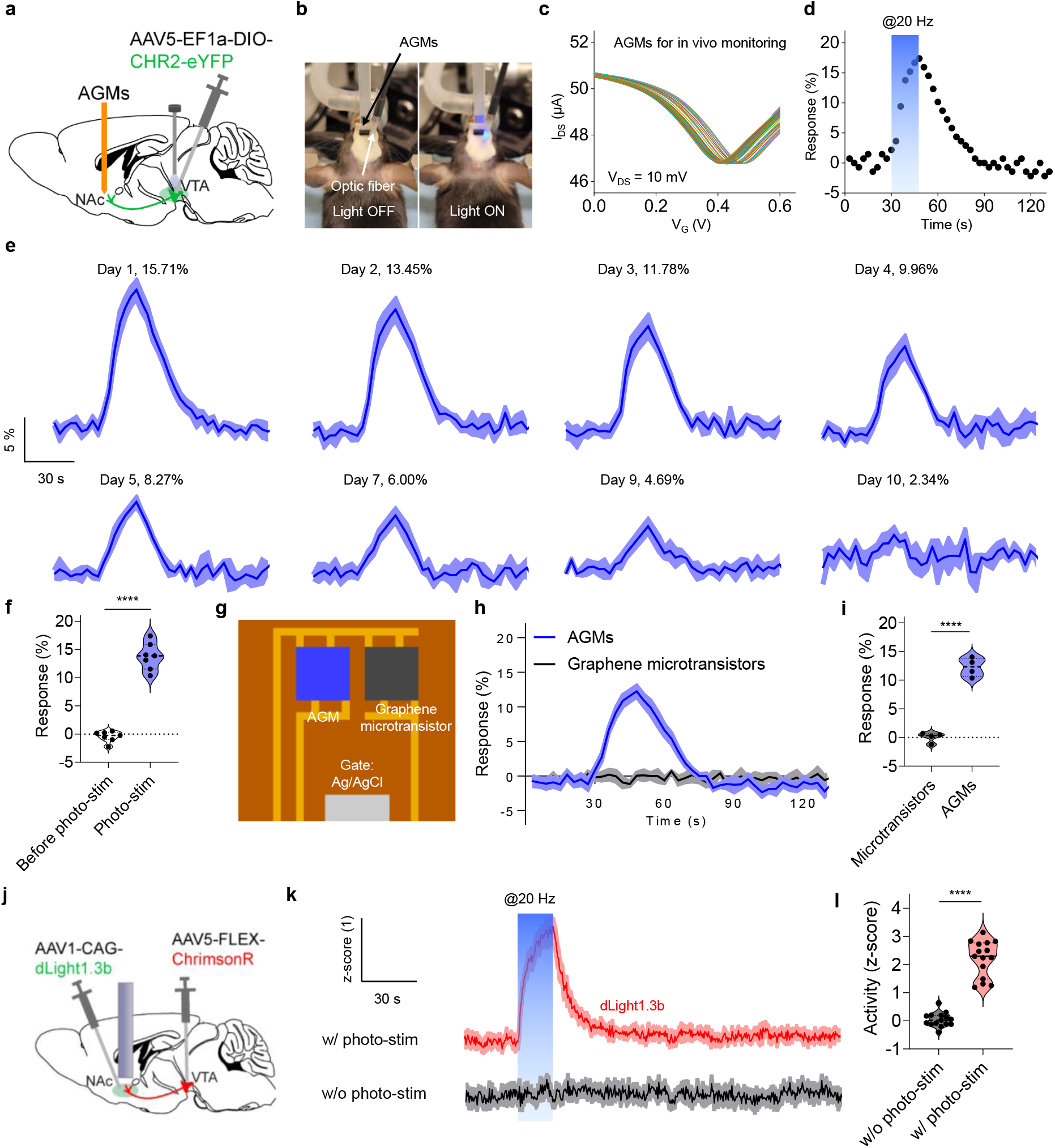
Long-term monitoring of optically evoked dopamine release *in vivo*. (**a**) Schematic illustration of AGMs implanted in NAcSh with ChR2 injected and optic fiber implanted in VTA. (**b**) Optical images of a mouse implanted with AGMs and optical fiber. (**c**) Representative transfer curves of the AGMs during photostimulation process. The time interval between each transfer curve is 3 s, and the scanning step is 3 mV for V_G_. (**d**) The sensor response to dopamine release evoked by photostimulation at 20 Hz and 5 ms pulse width. (**e**) Long-term monitoring of optogenetically evoked dopamine release *in vivo* using AGMs with surface coatings. The shading area represents ± SEM from three samples. (**f**) Comparison of the sensor response to dopamine release before and during photostimulation on day 1. n = 7; \*\*\*\**P* < 0.0001. All data are represented as means ± SEM. (**g**) Schematic illustration of an implantable probe that contains graphene microtransistor with and without the aptamer functionalization. (**h**) Simultaneous operation of graphene microtransistor and AGM channels for real-time dopamine monitoring during photostimulation. The shading area represents ± SEM from four samples. (**i**) Sensor response to optogenetically evoked dopamine release for graphene microtransistor and AGM channels. n = 4; \*\*\*\**P* < 0.0001. All data are represented as means ± SEM. (**j**) Schematic illustration of the mouse brain injected with ChrimsonR in VTA, and dLight injected and optic fiber implanted in NAcSh. (**k**) Averaged traces of dLight1.3b response in NAcSh with and without the photostimulation (20 Hz, 5 ms pulse width). The shading area represents ± SEM from 5 mice (15 tests). (**l**) Calculated response to dopamine release detected by dLight1.3b. n = 15; \*\*\*\**P* < 0.0001. All data are represented as means ± SEM.

To further determine whether the sensor response is due to the interaction between the optically-induced dopamine release and the aptamer monolayer on the sensor surface, we designed an implantable neural probe that contains both AGM channel and graphene microtransistor only channel (**Fig. 5g**). The probe was then passivated with pyrene-PEG5-alcohol and coated with rutin- doped polyacrylamide hydrogel. The probe with these two channels was implanted into the NAcSh of DAT-cre mice to monitor optically evoked dopamine release. More specifically, when the light was delivered through VTA terminal stimulation (20 Hz, 5 ms pulse width), the two channels on the implanted probe were monitored simultaneously. **Fig. 5h** shows a rapid increase in the sensor response for the AGM channel, while the graphene microtransistor channel (without aptamer functionalization) shows no clear response, indicating that the sensor response is attributed to the interaction between the surface functionalized aptamers and released dopamine molecules (**Fig. 5i**). Similarly, the AGM channel exhibits clear responses to dopamine release over one week whereas the graphene microtransistor channel shows a negligible response (**Supplementary Fig. 10**). We reason that the hydrogel coating could eliminate the influence from the electrophysiological signal through the gate electrode. Moreover, the dopamine release signal was recorded every 3 s, whereas the temporal resolution of the electrophysiological signal is in the range of milliseconds. These studies further support that the obtained transfer curve shift is indeed due to the dopamine release induced by photostimulation. There is negligible effect of the photostimulation on sensor response in control mice that were injected with a control ChR2-absent vector, AAV5-EF1a-DIO-eYFP, in VTA (**Supplementary Fig. 11**), indicating that the sensor response was generated by optically-evoked dopamine release in the NAcSh rather than photoartifact of the photostimulation. Overall, these results indicate that DNase inhibitor-doped hydrogel coatings can effectively extend the lifetime of AGMs to over one week for *in vivo* monitoring of neurotransmitter release.

To compare the AGM biosensor with other emerging biosensors, genetically encoded fluorescent indicators, we injected Cre-recombinase-dependent viral vectors containing the light-sensitive cation channel ChrimsonR into the VTA of DAT-cre mice, along with genetically encoded fluorescent dopamine indicator dLight^56^ and a fiber photometry implant into the NAcSh (**Fig. 5j**). We then delivered photostimulation (20 Hz, 5 ms pulse width) above the NAcSh. A significant peak response was obtained from stimulation followed by a decay in signal post-stimulation, similar to the response of the AGMs (**Figs. 5k and l**). These results indicate that the hardware-based AGMs demonstrate comparable performance for dopamine release detection to comparable and existing sensing technology. The genetically encoded fluorescent indicator offers a high-resolution benchmark for our AGMs. This technology, however, has limited translational potential due to complicated genetic modifications.

### *In vivo* dopamine monitoring and longevity for AGMs in freely moving behaviors

We next measured the capability of AGMs to record *in vivo* endogenous dopamine release induced through behaviors such as righting reflex and sucrose reward consumption. RORR is an established behavioral means to assess behavioral responsiveness in rodents, and it has been reported that NAcSh dopamine increases during this process^57^. RORR is assessed as the time for rodent take to return to a prone position following anesthesia. We used this method to establish the ability of our device with surface coating to monitor endogenous dopamine dynamics. As shown in **Figs. 6a** and **b**, the RORR process is demonstrated on a mouse following AGM implantation into the VTA. Eight days following implantation, transfer curves were continuously monitored at the time of righting, and a rapid increase in sensor response was obtained by the AGMs compared to baseline measurements of the transfer curves (**Fig. 6c)**. We then calculated the overall response before and during RORR and found a significant difference between the baseline and when the mice demonstrate RORR, suggesting the dopamine release in the NAcSh during this process (**Fig. 6d**).

**Figure 6.**
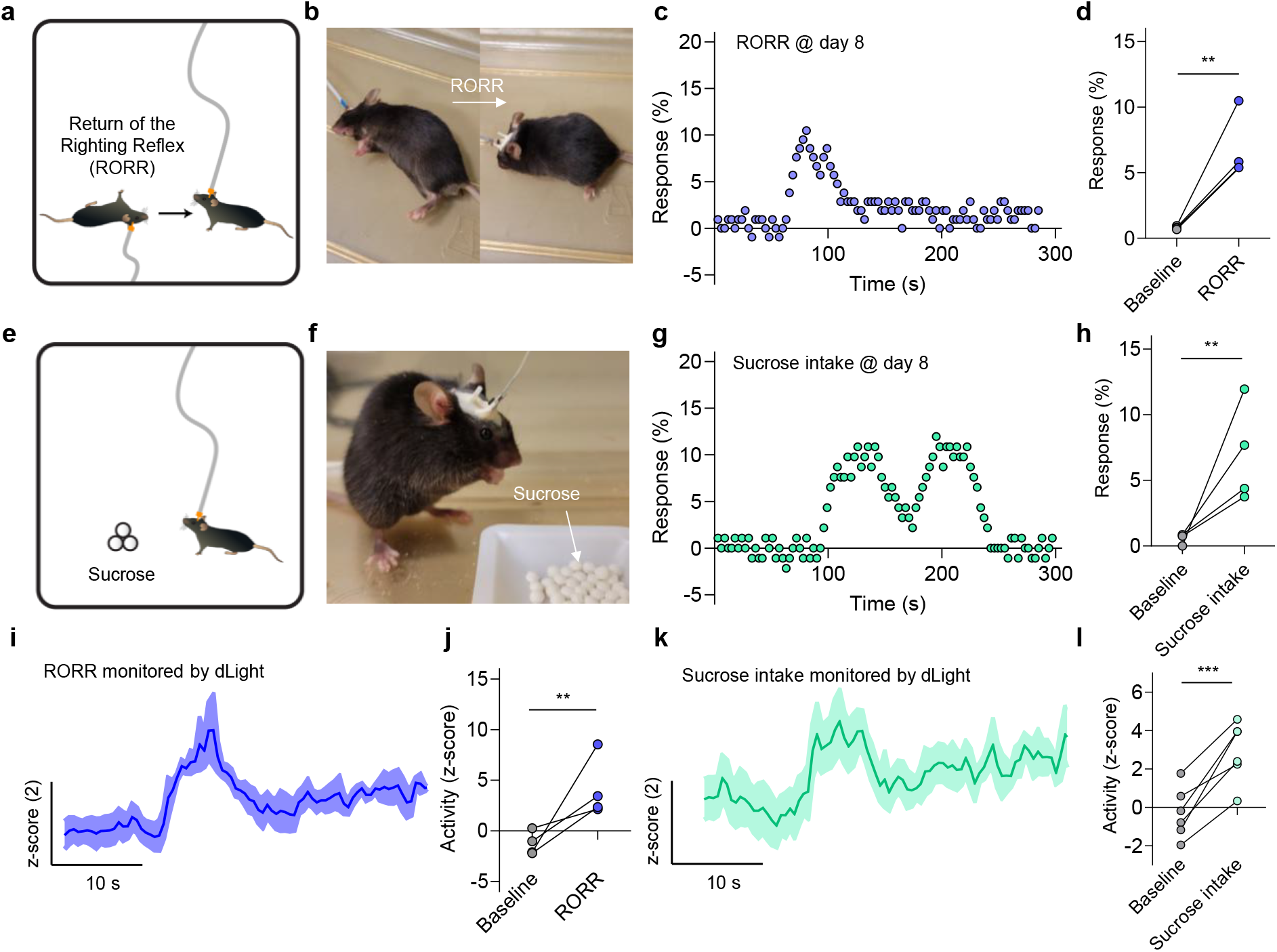
Long-term monitoring of dopamine release during animal behavior events. (**a**) Schematic illustration of the return of righting reflex (RORR). (**b**) Optical images of a mouse during RORR. (**c**) The representative sensor response during RORR after eight days of sensor implantation. The time interval between each data point is 3 s. (**d**) Comparison of the sensor response before and during the RORR. n = 4; \*\**P* < 0.01. All data are represented as means ± SEM. (**e**) Schematic illustration of sucrose intake behavior. (**f**) Image of a mouse during sucrose intake behavior. (**g**) The representative sensor response during sucrose intake after eight days of sensor implantation. (**h**) Comparison of the sensor response before and during sucrose intake. n = 4; \*\**P* < 0.01. All data are represented as means ± SEM. (**i**) Representative trances of dopamine release signal during RORR detected by dLight method. (**j**) Comparison of the dLight signal response to dopamine release during the RORR. n = 4, \*\**P* < 0.01. All data are represented as means ± SEM. (**k**) Representative trances of dopamine release signal during sucrose intake detected by dLight method. (**l**) Comparison of the dLight signal response to dopamine during the sucrose intake. n = 6, \*\*\**P* < 0.001. All data are represented as means ± SEM.

We also tested the capabilities of AGMs with sucrose reward consumption, which is well-established to evoke robust dopamine release within the NAcSh^58–60^. After 8 days of sensor implantation in the NAcSh, mice were placed into a chamber with free access to sucrose pellets while continuous dopamine recording was collected (**Figs. 6e** and **f**). An increase in dopamine signal response was observed when the mice began consuming sucrose pellets, and the signal gradually returned to baseline when the mice stop feeding (**Figs. 6g** and **h**). When comparing baseline measurements to the peak response during sucrose reward consumption, a significant difference was observed, establishing detection of behaviorally-evoked endogenous dopamine release by our AGM-coating strategy. Comparable dopamine release was also detected by the dLight biosensor in a series of side-by-side experiments, further establishing that the AGMs can be reliably used to monitor dopamine *in vivo* in freely moving animals (**Figs. 6i-l**). These successful sub-chronic monitoring of behaviorally-induced *in vivo* dopamine release further highlights the feasibility of surface-coated AGMs for widespread use in behavioral studies and neural circuit dissection over long time frames and in more complex scenarios.

## Discussion

Aptamer-based electrochemical biosensors are emerging high-performance sensing platforms for real-time molecular monitoring in undiluted whole blood samples and complex *in vivo* biological environments. Yet, the longevity of these sensors has been a significant challenge. To date, the *in vivo* operation of existing aptamer-based electrochemical biosensors is limited to a few hours (12 hours at most), which greatly hampers applications in which one to several weeks-long operation is needed, like therapeutic drug monitoring, neurochemical monitoring, and protein detection for immunoengineering. Here, we demonstrate significant progress in solving this long-standing challenge by developing a first-generation long-term, real-time molecular monitoring platform, AGMs, in undiluted whole blood samples and complex *in vivo* environments. The long-term operation is achieved by a combined effect of: 1) pyrene-PEG5-alcohol passivation and 2) DNase inhibitor-doped polyacrylamide hydrogel coatings. MTT analysis of forebrain organoids cocultured with these coatings and *in vivo* immunohistochemistry study indicate their biocompatibility for use in *in vivo* real-time molecular monitoring. The reported sensors allow for long-term (over one week) monitoring of optically evoked dopamine release *in vivo* using mice models. In addition, the monitoring of animal behavior (RORR and sucrose reward consumption) induced dopamine release by the AGMs is successfully demonstrated. Compared with the hour-long operation of existing aptamer-based biosensors in *in vivo* studies, this multi-day (over one week) operation significantly extends the *in vivo* operation lifetime and marks a new benchmark for aptamer-based biosensors in complex *in vivo* environments.

In addition to the significantly improved *in vivo* operational lifetime, the AGMs simultaneously transduce the interaction between aptamers and targets of interest and amplify the resulting signals, which makes it possible for the miniaturization of sensor size down to micrometer scale (50 μm × 50 μm of each sensor) without sacrificing the signal-to-noise ratio. In contrast, when the electrode size of aptamer-electrochemical biosensors is miniaturized down to such micrometer scale without incorporating porous structures, the magnitude of the resulting current would be as low as nA. The reported minimum current range for the state-of-the-art miniaturized potentiostat module (EmStat Pico) is 100 nA. Therefore, the accurate detection of such low currents often requires sophisticated and bulky electrochemical data acquisition systems, which adds additional challenges to customized circuit design for developing sensors for real-time molecular monitoring in a wearable platform like CGM.

A recent study systematically assessed the degradation mechanisms of self-assembled monolayer of aptamer-electrochemical biosensors and expected one week-long operation in bovine serum at 37 °C^61^, yet our study pushes these limits in several ways: (1) only 3 days of operation in bovine serum at 37 °C is experimentally demonstrated in this recent study. In contrast, we reported the longer operation of our AGMs in more complex undiluted whole blood samples (over 11 days) instead of simple serums. (2) we demonstrated the multi-day time scale (over one week) operation of our AGMs *in vivo*.

Overall, the surface-coated AGMs extend the *in vivo* operation lifetime of aptamer-based biosensors from a few hours to week-long time scales, which opens the avenue for long-term molecular sensing in complex biological environments in several settings. These include behavioral neural circuit analysis, disease modeling, therapeutic drug monitoring, immune monitoring, and chronic disease management. Future work includes further extending the lifetime of AGMs to even longer time scales (like CGM) by optimizing formulations of surface coatings, developing platforms for multiplexed monitoring of neurotransmitters, alongside neuropeptides, in freely moving animals, and developing a feedback-controlled system for molecular monitoring and on-demand medical interventions. The technology we present here represents a first-generation long-term aptamer-based biosensor for *in vivo* molecular monitoring and serves as a unique template for the design of future iterations with added functionality, longevity, and operational modes.

## Methods Materials

Chemical vapor deposition graphene sample was purchased from Graphenea. Dopamine, tetrabutylammonium hexafluorophosphate, p-phenylenediamine, N-ethyl-N ′ -(3-(dimethylamino)propyl)carbodiimide (EDC), N-hydroxysuccinimide (NHS), pyrene-PEG5- alcohol, N,N-diethylacrylamide, N-hydroxyethyl acrylamide, lithium phenyl-2,4,6- trimethylbenzoylphosphinate (LAP), N,N-methylenebis(acrylamide), and bovine serum albumin (BSA) were purchased from Sigma Aldrich and used as received. Artificial cerebrospinal fluid (aCSF) was purchased from EcocyteShop and used following the instructions. Rat CSF and EDTA-treated rat whole blood were obtained from BioIVT. The customized dopamine and serotonin aptamers were purchased from Integrated DNA Technologies. Polyimide film (DuPont™ Kapton®) was purchased from American Durafilm Co., Inc. Liquid polyimide (PI2545) was purchased from HD MicroSystems™.

### Fabrication of graphene microtransistor devices

Graphene microtransistor devices were prepared using the photolithography method, as described previously^35, 36, 38^. Briefly, polyimide film (76 μm thick) was laminated on a glass slide with a Polydimethylsiloxane (PDMS) adhesion layer. The liquid PI2545 was spin-coated onto the polyimide film and cured in a vacuum oven to form an additional clean and uniform polyimide layer (5 μm thick). The source, drain, and onboard gate terminals were formed by directly writing on the surface of the polyimide substrate coated with AZ 5214 using a maskless aligner (μMLA, Heidelberg Instruments), followed by the metal deposition (15 nm Cr/90 nm Au) and lift-off processes. Another UV exposure, metal deposition, and lift-off were performed to define the Ag-based gate terminal. Finally, a small droplet of diluted bleach solution (sodium hypochlorite, 5 μL) was added to the Ag electrode surface, resulting in Ag/AgCl gate electrode. Chemical vapor deposition graphene on copper was then transferred and patterned to bridge the source and drain electrodes. SU-8 (800 nm thick) was used to encapsulate the electrodes to prevent possible leakage during the test, resulting in an active sensing area of graphene being 50 µm × 50 µm.

### Aptamer functionalization using the covalent method

The amino group was grafted onto the graphene surface based on the electrochemical grafting method^36^. Here, NaNO_2_ solution (10 mL, 4 mM) was first added into HCl solution (1 M) containing 10 mM phenylenediamine (PPD). After nitrogen degassing for 5 min, diazonium salt (ClN_2_^+^-Ph-NH_2_) was formed by putting the resulting solution in an ice-water bath for 10 min. To graft the amino group onto graphene surface, five cycles of cyclic voltammetry (CV) scans were performed. The scan rate was 100 mV/s and the potential window was set from-0.6 to 0.5 V. After that, the graphene microtransistors were sequentially rinsed with acetonitrile and ultrapure DI water. The obtained amino group grafted graphene microtransistors were then incubated with 6 mM EDC (N-ethyl-N ′ -(3-(dimethylamino)propyl)carbodiimide), 3 mM NHS (N-hydroxysuccinimide), and 3 μM carboxyl group-modified dopamine aptamer (5′/COOH/CGA CGC CAG TTT GAA GGT TCG TTC GCA GGT GTG GAG TGA CGT CG) in 1× PBS buffer at room temperature for 2 h, followed by washing with aCSF buffer and blowing dry with N_2_ gas.

### Aptamer functionalization using the non-covalent method

A pyrene molecule was added to the dopamine aptamer during the synthesis process by Integrated DNA Technologies (IDT). The as-received pyrene-tagged dopamine aptamer (5’/pyrene/CGA CGC CAG TTT GAA GGT TCG TTC GCA GGT GTG GAG TGA CGT CG-3’) was diluted to its working concentration (1 µM) with aCSF solution. The dopamine aptamer solution was then heated under 85-95 °C in the water bath for 5 minutes and slowly cooled down to room temperature. Finally, the graphene microtransistors were incubated with the prepared solution for 2 hours at room temperature to functionalize the graphene surface with aptamers, followed by washing with aCSF buffer and drying with N_2_ gas. For AGM serotonin sensors, a pyrene linker (1-pyrenebutyric acid N-hydroxysuccinimide ester, 5 mM in methanol) was added to graphene surface for 4 hours at room temperature, followed by the incubation with 1 µM serotonin aptamer (5’/AmC6/CGA CTG GTA GGC AGA TAG GGG AAG CTG ATT CGA TGC GTG GGT CG-3’) at room temperature for 2 hours. The obtained AGM serotonin sensors were washed with fresh aCSF buffer and dried with N_2_ gas for later use.

### Surface passivation with pyrene-PEG5-alcohol

The pyrene-PEG5-alcohol powder was dissolved in DI water to prepare a concentration of 1 µM pyrene-PEG5-alcohol solution. Then the dopamine aptamer-functionalized graphene microtransisotrs were incubated in the prepared solution at room temperature for 2 hours. The resulting AGMs with pyrene-PEG5-alcohol passivation were finally rinsed with aCSF solution.

### Preparation of polyacrylamide hydrogels

The synthesis of polyacrylamide hydrogel started with preparing the prepolymer aqueous solution, which contains hydroxyethylacrylamide (monomer, 10 wt%), diethylacrylamide (monomer, 10 wt%), lithium phenyl-2,4,6-trimethylbenzoylphosphinate (photoinitiator, 1 wt%), and N, N′-methylenebisacrylamide (crosslinker, 1 wt%). The AGMs were then immersed into the monomer solution for 5 s and repeated for 3 times. The prepolymer solution-coated sensors were finally exposed to UV light (365 nm) for 3 minutes. The AGMs with pyrene-PEG5-alcohol and polyacrylamide hydrogel coatings were kept in aCSF solution for later use.

### Surface coating with rutin-doped polyacrylamide hydrogels

For sensors coated with rutin-doped polyacrylamide hydrogel, the prepolymer aqueous solution consists of hydroxyethylacrylamide solution (10 wt%), diethylacrylamide (10 wt%), lithium phenyl-2,4,6-trimethylbenzoylphosphinate (1 wt%) and N, N’-methylenebisacrylamide (crosslinker, 1 wt%) was mixed with rutin (150 µM). Then the sensors were coated with the prepared solution for 5 s and repeated for 3 times, followed by illumination with 365 nm UV light for 3 minutes. The resulting sensors were kept in aCSF solution for later use.

### Cell viability test

#### Maintenance of human induced pluripotent stem cell (hiPSC) culture

Human iPSK3 cells were maintained in mTeSR plus serum-free medium (StemCell Technologies, Inc., Vancouver, Canada) on a growth factor reduced Matrigel-coated surface (Corning Incorporated, Corning, NY). Cells were passaged every 4-6 days using Accutase and seeded at a density of 1 × 10^6^ cells per six-well plate in the presence of rho-associated protein kinase (ROCK) inhibitor Y27632 (10 μM, Sigma) for the first 24 hours.

#### Forebrain organoid differentiation

Human iPSK3 cells were seeded at a density of 3 × 10^5^ cells per well into 24 well, ultra-low attachment plates (Corning Incorporated) in differentiation medium composed of Dulbecco’s Modified Eagle Medium/ Nutrient Mixture F-12 (DMEM/F-12) with 2% B-27 serum-free supplement (Life Technologies, Carlsbad, CA). Y27632 (10 μM, Sigma) was included in the differentiation medium for the first 24 hours. After 24 hours, the Y27632 was removed and the formed embryoid bodies were treated with the dual SMAD inhibitors SB431542 (10 μM, Sigma-Aldrich, St. Louis, MO) and LDN193189 (100 μM, Sigma) for 7 days. On day 8, cells were treated with 10 ng/mL fibroblast growth factor (FGF)-2 (Life Technologies) for another 7 days. After day 15, the spheroids were cultured in DMEM/F-12 without any growth factors for 8 days. On day 23, spheroids were seeded onto a Matrigel-coated 96 well plate and allowed to attach for 2 days. Media changes were performed every 2 days.

### (4,5-dimethylthiazol-2-yl)-2,5-diphenyltetrazolium bromide (MTT) Assay

On day 25 of spheroid culture, AGMs with or without pyrene-PEG5-alcohol and rutin-doped hydrogel coatings were introduced to the culture. Cells were treated with a 0.5 mg/mL of MTT reagent (Sigma) after 1, 4, 8, 12, or 16 days of culture with the above-mentioned AGMs with or without surface passivation. The media containing MTT reagent was removed from cells. The formazan crystals were then washed with dimethyl sulfoxide (DMSO) and centrifuged at 800 g for 5 minutes. The absorbance of the supernatant was measured at 490 nm on a microplate reader (BioRad Laboratories, Hercules, CA).

### Animals

Adult DAT-IRES-Cre mice were used in this study. Mice were 8-10 weeks old at the time of test, weighing 20-30 g each, group-housed, given access to food pellets and water ad libitum, and maintained on a 12:12-hour light/dark cycle (lights on at 7:00 a.m.). Animals were held in a sound attenuated holding room facility in the lab starting at least one week prior to surgery, as well as post-surgery and throughout the duration of behavioral assays to minimize stress from transportation and disruption from foot traffic. All experimental procedures were approved by the Animal Care and Use Committee of the University of Washington and conformed to NIH guidelines.

### Stereotaxic surgery

All surgeries were performed under 4% isoflurane anesthesia (Piramal Healthcare, Maharashtra, India). For testing of AGMs, adult mice were injected unilaterally with 600 nL of either AAV5- EF1a-DIO-ChR2-eYFP virus (WUSTL Hope Center Viral Core, St. Louis, MO) or AAV5-EF1a- DIO-eYFP (WUSTL Hope Center Viral Core, St. Louis, MO) into the ventral tegmental area (AP: -3.25, ML: 0.5, DV: -4.4) using a Hamilton syringe with a blunted needle. Four to five weeks after virus injection, fiber optic ferrules were chronically implanted above the VTA (AP: -3.25, ML: 1.6, DV: -4.3 at 15° angle) and dental cement (Lang Dental, Wheeling, IL) was applied to hold the ferrules in place. The AGM probe was then implanted in the nucleus accumbens shell (AP: 1.45, ML: 0.65, DV: -4.5). For fiber photometry experiments using dLight, adult mice were injected unilaterally with 600 nL of AAV-CAG-dLight1.3b (Lin Tian Lab, UC Davis) into the NAcSh (AP: 1.45, ML: 0.65, DV: -4.5) using a Hamilton syringe with a blunted needle. Four to five weeks after virus injection, fiber photometry ferrules were chronically implanted into the NAcSh (AP: 1.45, ML: 0.65, DV: -4.5). Mice were allowed to recover for at least six weeks following infusion of the virus prior to further behavioral testing to ensure optimal viral expression and implant placement location. All experimental procedures were approved by the Animal Care and Use Committee of University of Washington and conformed to NIH guidelines.

### Open-field test (OFT)

OFT testing was performed in a custom-made white box (50 cm × 50 cm) within a sound-attenuated room maintained at 23°C. The ‘center’ area was defined as a square of 50% of the total OFT area. A light bulb mounted above provided 45 lux illumination, measured in the center of the arena. Mice were handled for 5 minutes per day for a week prior to the OFT testing. On the day of OFT testing, mice were habituated in a holding room for 30 minutes and then they were allowed to explore the entire chamber freely for 25 min. Sessions were recorded via a digital camera, and videos were analyzed offline using Ethovision XT 8.5 (Noldus Information Technologies). The open field was cleaned with 70% ethanol between each trial. Ethovision XT 8.5 (Noldus Information Technologies) quantified the total distance and time spent in the central and peripheral areas. Time spent in the center of the open field was used as a measure of anxiolysis. All experimental procedures were approved by the Animal Care and Use Committee of the University of Washington and conformed to NIH guidelines.

### Optogenetics stimulation

Following stereotaxic surgery and implantation of the AGM probe in NAc, mice were connected to a 473 nm diode-pumped solid-state laser (OEM Laser Systems). Laser power was adjusted to obtain ∼15 mW transmittance into the brain, at 20 Hz frequency with 5 ms pulse for 20 s. All experimental procedures were approved by the Animal Care and Use Committee of the University of Washington and conformed to NIH guidelines.

### Sucrose intake

Behaviorally tested mice were food deprived (mice still had access to water) for 24 hours prior to testing. Then, animals were given access to ∼1 g of sucrose pellets and had free access to these pellets for approximately 15 minutes. After testing, food-deprived animals were returned to an ad libitum diet. Sessions were recorded via a digital camera, and videos were analyzed offline using Ethovision XT 8.5 (Noldus Information Technologies). All experimental procedures were approved by the Animal Care and Use Committee of the University of Washington and conformed to NIH guidelines.

### Righting reflex

Behaviorally tested mice were anesthetized in an induction box with 2.5% isoflurane (Piramal Healthcare, Maharashtra, India) for ∼5 minutes. Following this, mice were removed from anesthesia and placed into a recovery cage to exposure to the open air, until they exhibited righting reflex. After sufficient recovery time, the mice were returned to their home cage. Sessions were recorded via a digital camera, and videos were analyzed offline using Ethovision XT 8.5 (Noldus Information Technologies). All experimental procedures were approved by the Animal Care and Use Committee of the University of Washington and conformed to NIH guidelines.

### Brain tissue collection

DAT-cre mice were used in harvested brain tissue experiments. Mie were anesthetized with sodium pentobarbital and transcardially perfused with ice-cold PBS, followed by 4% phosphate-buffered paraformaldehyde. Brains were removed, postfixed overnight in 4% paraformaldehyde, and then saturated in 30% phosphate-buffered sucrose for 2-4 days at 4°C. All experimental procedures were approved by the Animal Care and Use Committee of the University of Washington and conformed to NIH guidelines.

### Immunohistochemistry

The AGM probe was implanted in the nucleus accumbens (AP: 1.45, ML: 0.65, DV: -4.5) for 1 and 6 weeks. To determine the immune response to the implanted devices, we stained tissue at both time points for astrocytes and activated microglia. The primary antibodies used were anti-GFAP guinea pig (SySy, 1:500 dilution) and anti-Iba1 rabbit (Abcam, 1:500 dilution), and the secondary antibodies used were Alexa Fluor 555 goat anti-guinea pig (Life Technologies, 1:1000 dilution) and Alexa Fluor 488 goat anti-rabbit (Invitrogen, 1:1000 dilution). We used the same confocal microscope settings to determine that the immune response for astrocytes and microglia at the 1- week time point was greater than that at the 6-week time point. All experimental procedures were approved by the Animal Care and Use Committee of the University of Washington and conformed to NIH guidelines.

## Acknowledgement

This work is supported by NIH BRAIN Initiative RF1NS118287 (to Y.Z.). This work made use of the ProtoLaser U4, which was funded by the Defense University Research Instrumentation Program from the Office of Naval Research (N00014-21-1-2223).

## Author contributions

GW, ETZ, and YQ contributed equally to this work. GW, MRB, and YZ conceived of the idea and designed the experiments; GW and YQ designed and fabricated AGMs; ETZ and MRB performed *in vivo* studies in mice; CE, XC, and YL contributed to the cell culturing experiments; ZW and JH contributed to the synthesis of polyacrylamide hydrogels; YS performed the electrografting experiments; HL and ZW performed Raman characterizations; YS and HS contributed to the SEM characterizations; MJS contributed to the *in vivo* mice studies, XZ and SS contributed to the signal processing, RL contributed to the surface functionalization of aptamers; GW, ETZ, MRB, and YZ analyzed the data and wrote the manuscript. All authors discussed the results and contributed to the final manuscript.

## Statistical analysis

Statistical analysis was performed using GraphPad Prism 10. The number of animals used for each experiment is reported along with the sample size (n). Data from any failed devices were excluded from the analysis. All experimental results are presented as means ± standard error of the mean (SEM). For two-group comparisons, statistical significance was evaluated using one-tailed Student’s t tests. Multiple-group comparisons were assessed using one-way ANOVA with Tukey’s multiple comparisons test. \**P* < 0.05, \*\**P* < 0.01, \*\*\**P* < 0.001, and \*\*\*\**P* < 0.0001 are considered statistically significant.

## Notes

### Competing Interest Statement

The authors have declared no competing interest.

