## Supplementary information for "Long-Term *In Vivo* Molecular Monitoring Using Aptamer-Graphene Microtransistors"

**Supplementary Table 1**

| Target | Detection method | Surface coating | Test environment | Test period | References |
|---|---|---|---|---|---|
| Serotonin | Aptamer FET | NA | *In vivo* in mouse brain | 20 min | Sci. Adv. 2021[1] |
| Phenylalanine | EAB | NA | *In vivo* in rat jugular vein | 1 hour | Anal. Chem. 2021[2] |
| Tobramycin | EAB | Polysulfone | *In vivo* in rat jugular vein | 4-12 hours | Proc. Natl. Acad. Sci. U. S. A. 2017[3] |
| Vancomycin | EAB | Zwitterionic polybetaine-based hydrogel | *Ex vivo* in undiluted bovine serum at 37 °C | 3 days (*Ex vivo*) | ACS Sens. 2023[4] |
| Kanamycin | EAB | Polyacrylamide hydrogel | *In vivo* in rat femoral vein | 3.5 hours | Adv. Mater. 2022[5] |
| Dopamine | AGM | Pyrene-PEG5-alcohol and rutin-doped polyacrylamide hydrogel | *In vivo* in mouse brain | Over one week | This work |

### Calculation of graphene microtransistor transconductance

The efficient modulation of the carrier mobility ($\mu$) by the gate voltage ($V_G$) can be explained by the following equation[6-8]:

$$\mu = \frac{\partial I_{DS}}{\partial V_G} \cdot \frac{l}{w} \cdot \frac{1}{C_{TG} \cdot V_{DS}}$$

where w and l are the width and length of the graphene channel, respectively; $C_{TG}$ is the total gate capacitance; $V_{DS}$ is the source-drain voltage; $\partial I_{DS}/\partial V_G$, also called transconductance ($g_m$) is defined as the derivative of the transfer curve. In this study, as shown in Figure 5a, the sensor transfer curve is scanned in the gate voltage operational window from 0 to 0.6 V, and the voltage interval between two data points is 0.003 V. Based on these, the calculated transconductance curve is shown in Fig. 5b, in which the red dash line indicates the maximum transconductance in the left branch of the transfer curve (hole, $g_{m\_h}$).

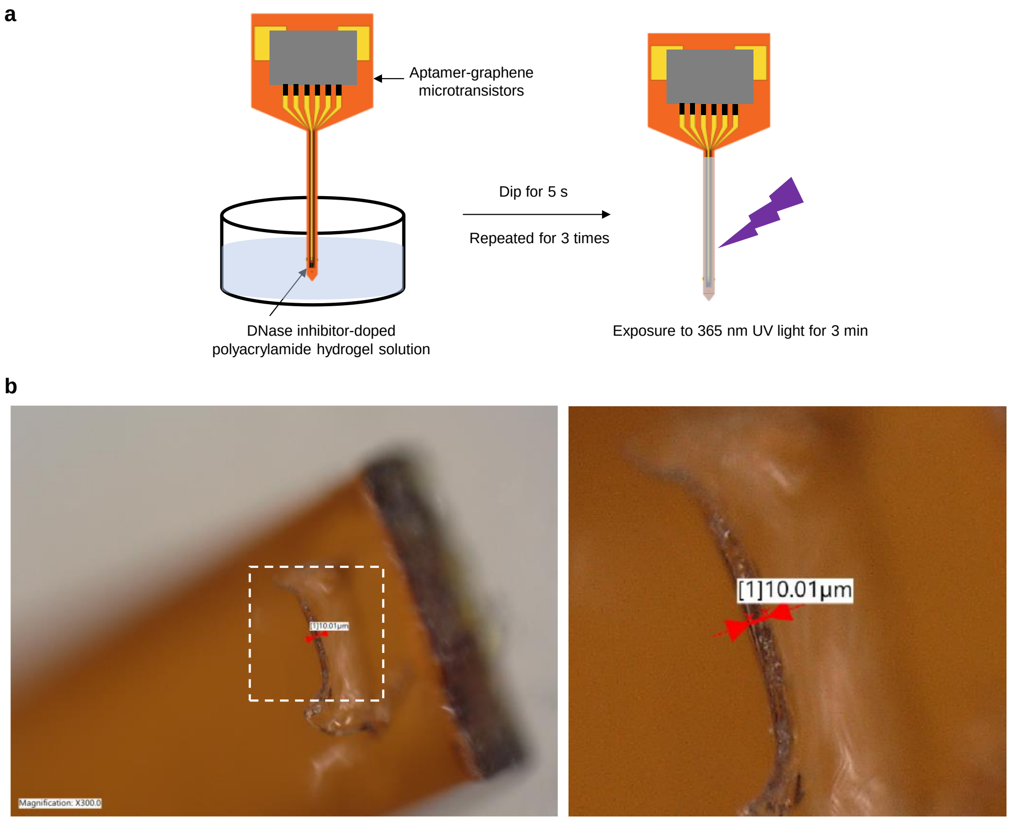


**Supplementary Fig. 1 | Surface coating with DNase inhibitor-doped polyacrylamide hydrogel.** **a**, General process of surface coating with DNase inhibitor-doped polyacrylamide hydrogel on the AGMs. **b**, Optical image shows the thickness of the polyacrylamide hydrogels coated on the surface of the AGMs.

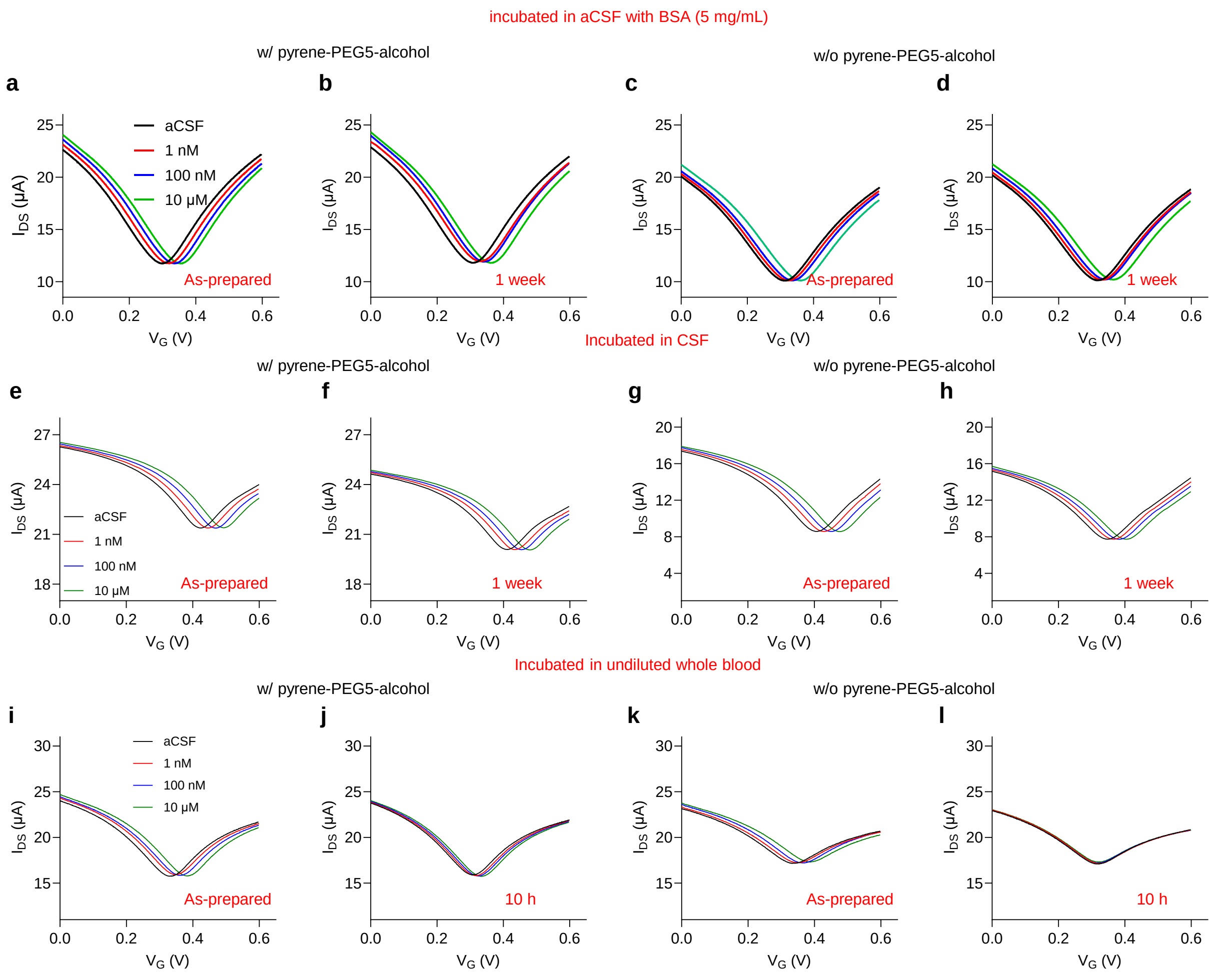


**Supplementary Fig. 2 | Long-term stability of pyrene-PEG5-alcohol passivated AGMs in aCSF, rat CSF, and undiluted whole blood samples at 37 ºC.** Transfer characteristics of surface passivated AGMs with pyrene-PEG5-alcohol (**a**) before and (**b**) after incubating in protein-rich aCSF solution (containing 5 mg/mL BSA protein) at 37 ºC for one week when exposed to various concentrations of dopamine. Transfer characteristics of AGMs without pyrene-PEG5-alcohol passivation (**c**) before and (**d**) after incubating in protein-rich aCSF solution (containing 5 mg/mL BSA protein) at 37 ºC for one week when exposed to various concentrations of dopamine. Transfer characteristics of surface passivated AGMs with pyrene-PEG5-alcohol (**e**) before and (**f**) after incubating in rat CSF at 37 ºC for one week when exposed to various concentrations of dopamine. Transfer characteristics of AGMs without pyrene-PEG5-alcohol (**g**) before and (**h**) after incubating in rat CSF at 37 ºC for one week when exposed to various concentrations of dopamine. Transfer characteristics of surface passivated AGMs with pyrene-PEG5-alcohol (**i**) before and (**j**) after incubating in undiluted rat whole blood for 10 hours when exposed to various concentrations of dopamine. Transfer characteristics of AGMs without pyrene-PEG5-alcohol (**k**) before and (**l**) after incubating in undiluted rat whole blood at 37 ºC for 10 hours when exposed to various concentrations of dopamine.

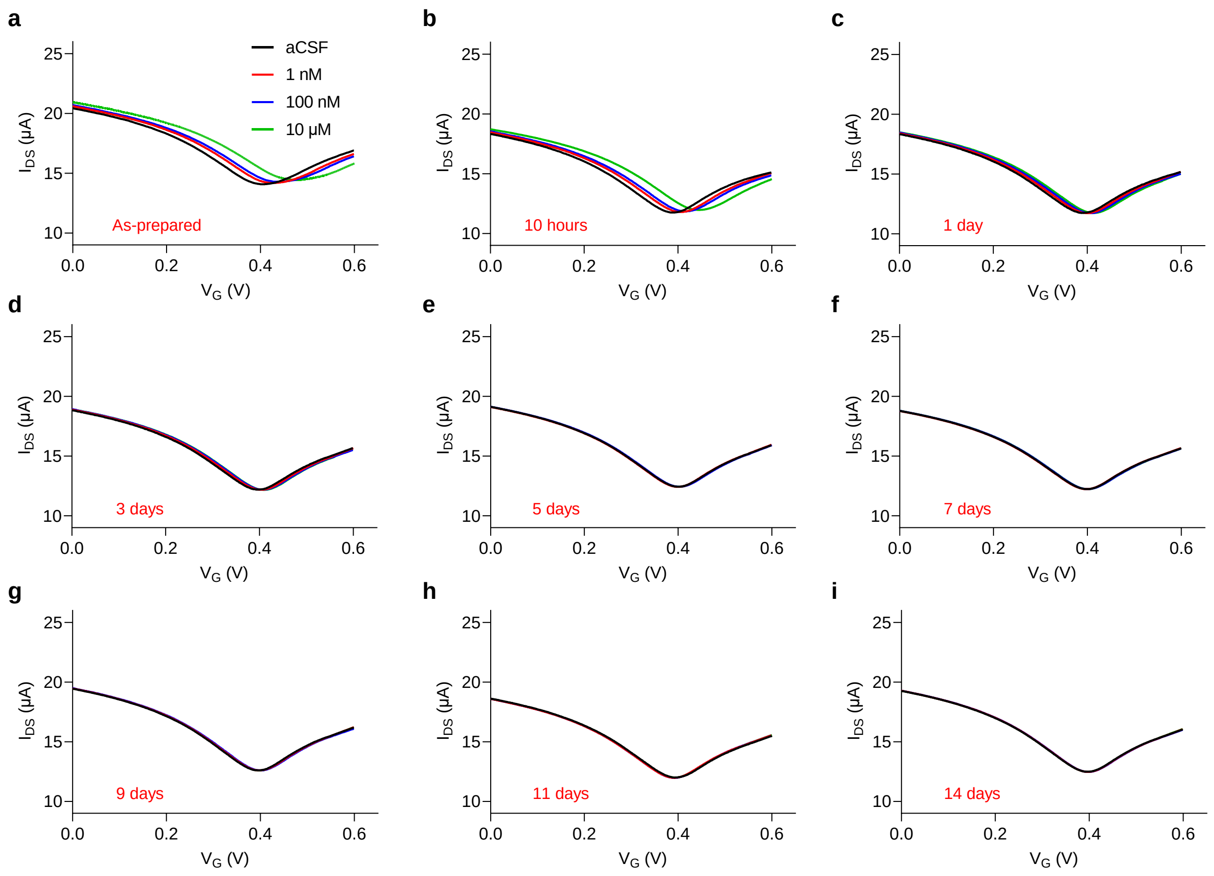


**Supplementary Fig. 3 | Long-term stability of the AGMs with pyrene-PEG5-alcohol and polyacrylamide hydrogel surface coatings in undiluted whole blood at 37 °C.** Transfer curves show the sensor response to different dopamine concentrations when the AGMs with pyrene-PEG5-alcohol and hydrogel coatings were incubated in undiluted blood sample for (**a**) 0 hour, (**b**) 10 hours, (**c**) 1 day, (**d**) 3 days, (**e**) 5 days, (**f**) 7 days, (**g**) 9 days, (**h**) 11 days, and (**i**) 14 days.

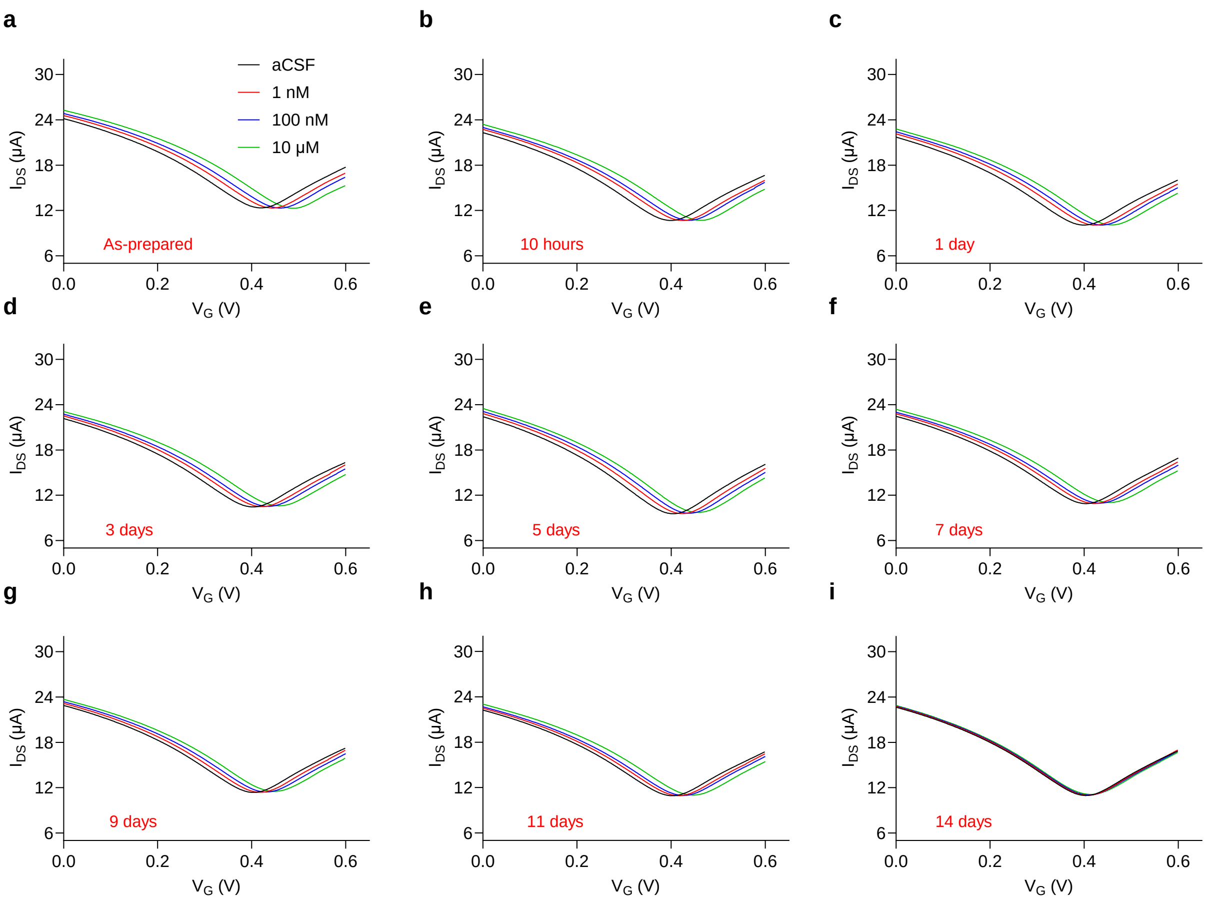


**Supplementary Fig. 4 | Long-term stability of the AGMs with pyrene-PEG5-alcohol and DNase inhibitor rutin-doped polyacrylamide hydrogel in undiluted whole blood samples at 37 ºC.** Transfer curves show the sensor response to different dopamine concentrations when the AGMs with pyrene-PEG5-alcohol and rutin-doped hydrogel coatings were incubated in undiluted blood sample for (**a**) 0 hour, (**b**) 10 hours, (**c**) 1 day, (**d**) 3 days, (**e**) 5 days, (**f**) 7 days, (**g**) 9 days, (**h**) 11 days, and (**i**) 14 days.

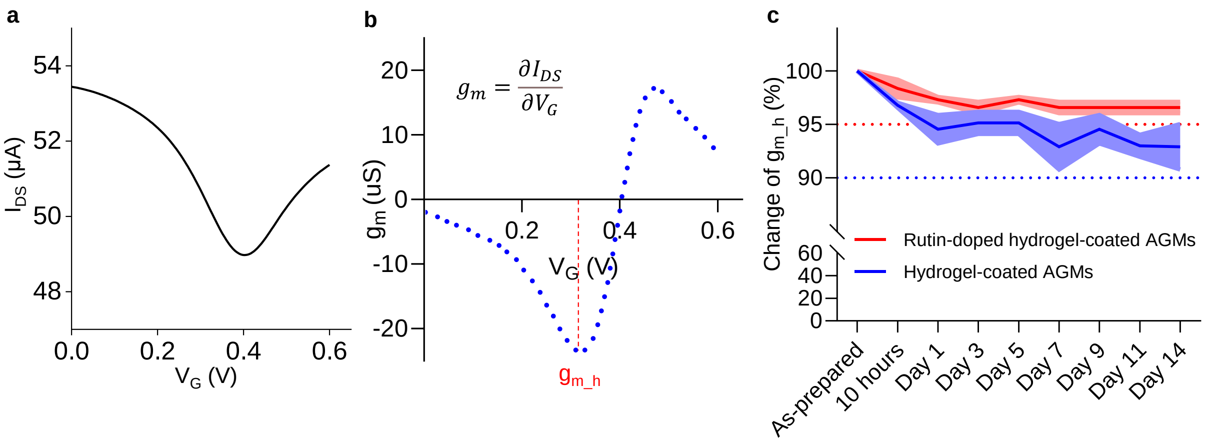


**Supplementary Fig. 5 | Transconductance calculations. a**, A representative transfer curve of the AGM. **b**, Derivative of the transfer curve in (**a**) to calculate the transconductance. **c**, The relative change of maximum transconductance ($g_{m\_h}$) of AGM with (red) and without (blue) rutin coating after incubating in undiluted blood for two weeks. The shading area represents ± SEM from three samples.

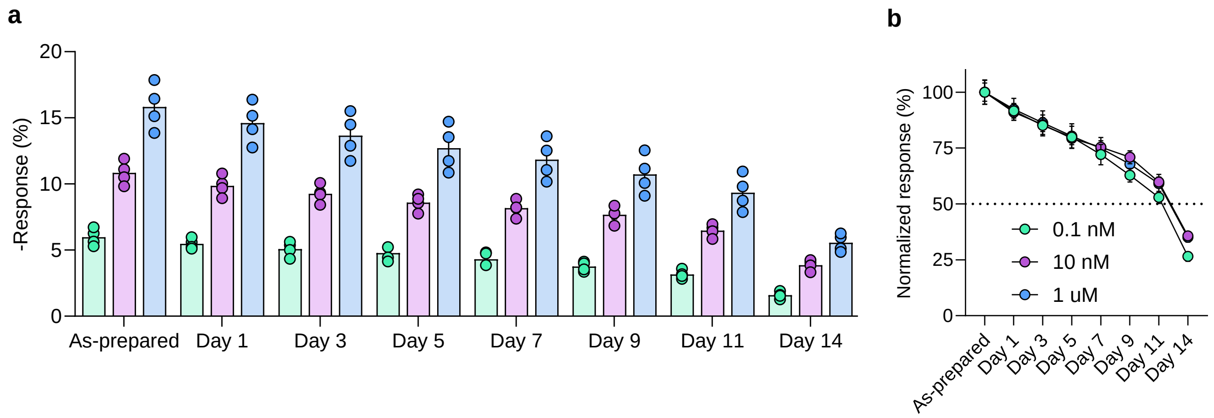


**Supplementary Fig. 6 | Long-term stability of AGM serotonin sensors with surface passivation in undiluted whole blood at 37 ºC. a**, The response of AGM serotonin sensors coated with pyrene-PEG5-alcohol and rutin-doped polyacrylamide hydrogel incubated in undiluted rat blood samples at 37 ºC. n = 4. **b**, Normalized sensor response after incubating in undiluted whole blood at different time points (1 day, 3 days, 5 days, 7 days, 9 days, 11 days, and 14 days) for pyrene-PEG5-alcohol and rutin-doped polyacrylamide hydrogel-coated AGM serotonin sensors. All data are represented as means ± SEM.

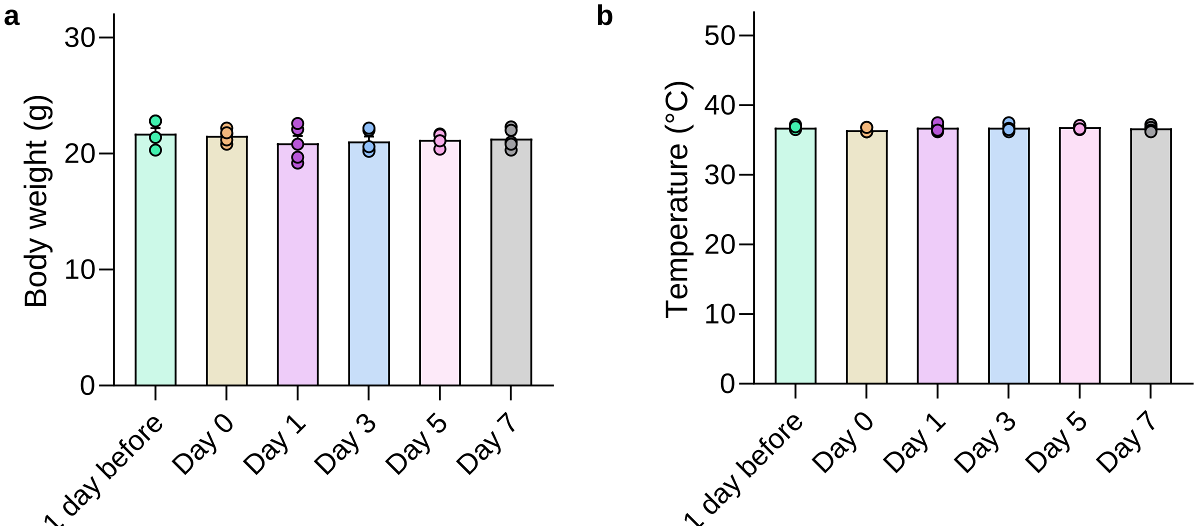


**Supplementary Fig. 7 | Impact of sensor implantation on animals. a**, Body weight of the mice as a function of time before and after implantation. **b**, Variation of body temperature of the mice before and after implantation. n = 5.

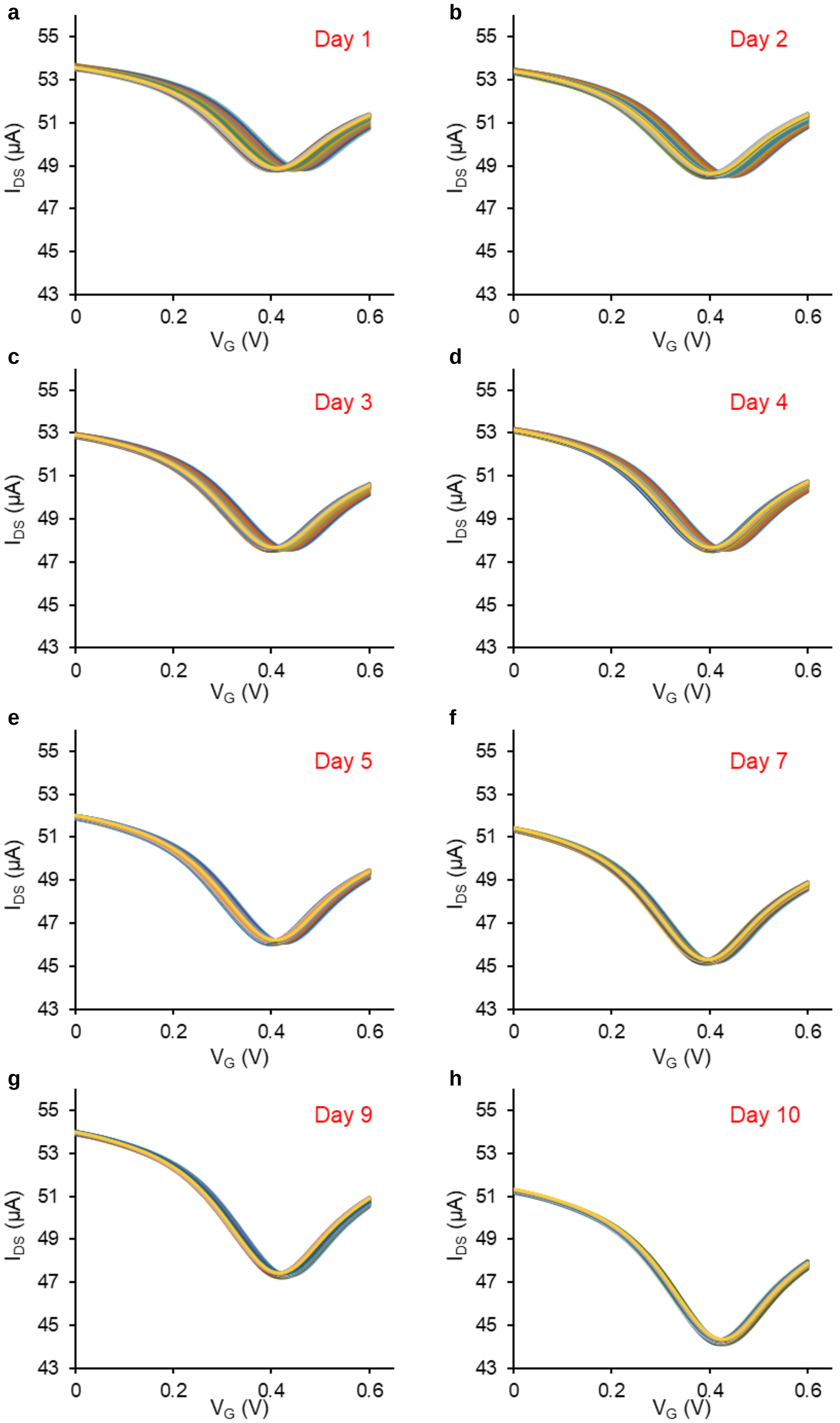


**Supplementary Fig. 8 | Sensor responses to dopamine release evoked by photostimulation.** Continuous transfer curve monitoring *in vivo* during photostimulation (20 Hz and 5 ms pulse width) after the AGMs with surface coatings were implanted into the NAcSh with ChR2 injected in VTA for (**a**) 1 day, (**b**) 2 days, (**c**) 3 days, (**d**) 4 days, (**e**) 5 days, (**f**) 7 days, (**g**) 9 days, and (**h**) 10 days. The time interval between each transfer curve is 3 s, and the scanning step is 3 mV for $V_G$.

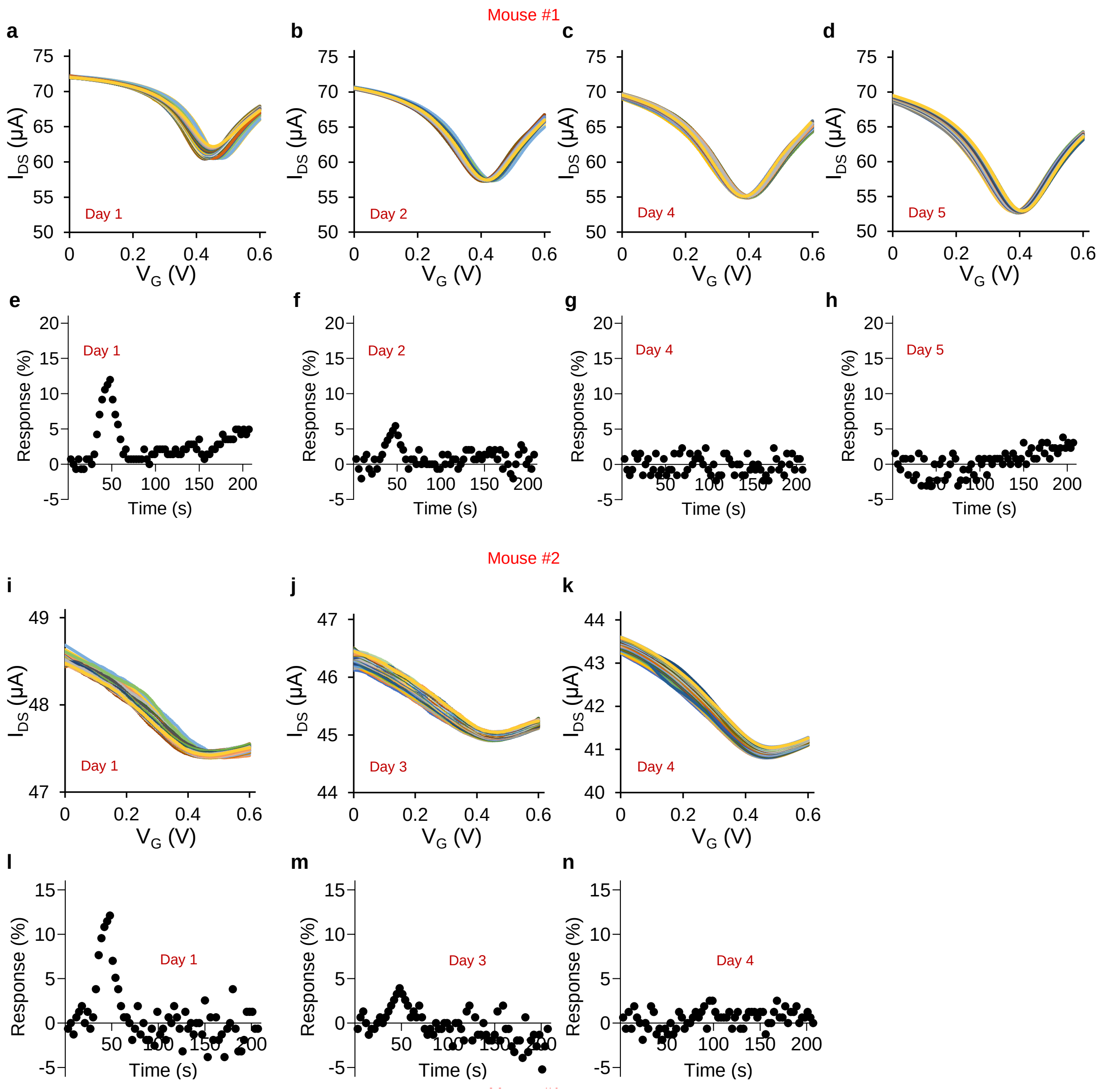


**Supplementary Fig. 9 | AGMs with pyrene-PEG5-alcohol and polyacrylamide hydrogel coatings for real-time dopamine monitoring during photostimulation.** Continuous transfer curve monitoring *in vivo* during photostimulation (20 Hz and 5 ms pulse width) after the AGMs with pyrene-PEG5-alcohol and hydrogel coatings were implanted into the NAcSh with ChR2 injected in VTA of mouse 1 for (**a**) 1 day, (**b**) 2 days, (**c**) 4 days, and (**d**) 5 days. Calculated response of the AGM and graphene microtransistor channels for *in vivo* monitoring of optically evoked dopamine release for (**e**) 1 day, (**f**) 2 days, (**g**) 4 days, and (**h**) 5 days. Continuous transfer curve monitoring *in vivo* during photostimulation (20 Hz and 5 ms pulse width) after the AGMs with pyrene-PEG5-alcohol and hydrogel coatings were implanted into the NAcSh with ChR2 injected in VTA of mouse 2 for (**i**) 1 day, (**j**) 3 days, and (**k**) 4 days. Calculated response of the AGM and graphene microtransistor channels for *in vivo* monitoring of optically evoked dopamine release for (**l**) 1 day, (**m**) 3 days, and (**n**) 4 days. The time interval between each transfer curve is 3 s and the scanning step is 3 mV for $V_G$.

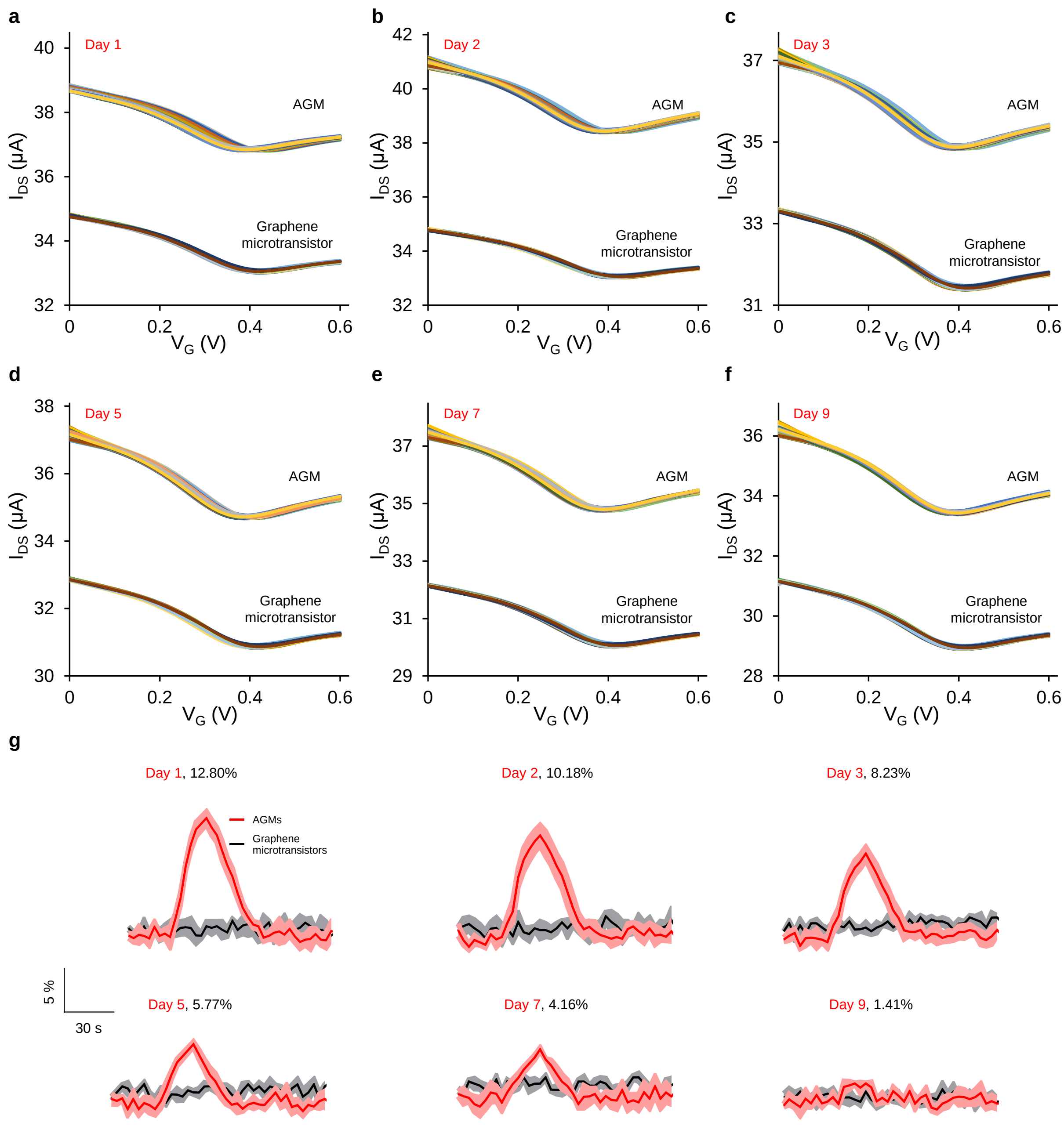


**Supplementary Fig. 10 | Simultaneous operation of graphene microtransistors and AGMs for real-time dopamine monitoring during photostimulation.** Continuous transfer curve monitoring *in vivo* during photostimulation (20 Hz and 5 ms pulse width) after the probe with AGM and graphene microtransistor channels with surface coatings was implanted into the NAcSh with ChR2 injected in VTA for (**a**) 1 day, (**b**) 2 days, (**c**) 3 days, (**d**) 5 days, (**e**) 7 days, and (**f**) 9 days. The time interval between each transfer curve is 3 s and the scanning step is 3 mV for $V_G$. (**g**) Overall response of AGM and graphene microtransistor channels to optically evoked dopamine release over 9 days. The shading area represents ± SEM from three samples.

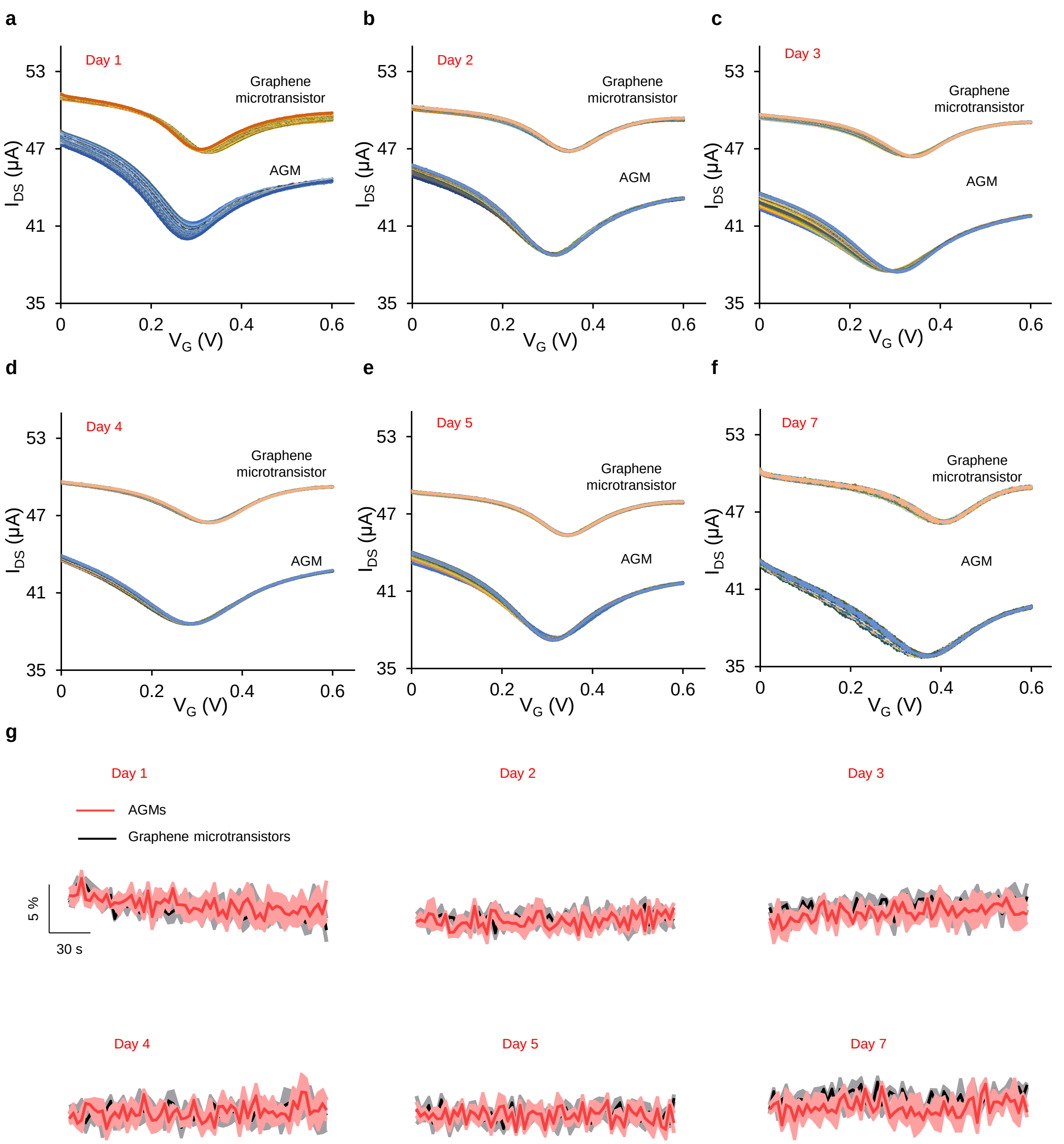


**Supplementary Fig. 11 | Sensor response in control mice injected with a control ChR2-absent vector, AAV5-EF1a-DIO-eYFP, in VTA.** Continuous transfer curve monitoring *in vivo* during photostimulation (20 Hz and 5 ms pulse width) after the probe with AGM and graphene microtransistor channels with surface coatings was implanted into the NAcSh with eYFP injected VTA for (**a**) 1 day, (**b**) 2 days, (**c**) 3 days, (**d**) 4 days, (**e**) 5 days, and (**f**) 7 days. (**g**) Overall response of the implantable neural probe with the AGM and graphene microtransistor channels to photostimulation evoked dopamine release after implantation over 7 days. The shading area represents ± SEM from three samples.
